# Heterotypic Assembly Mechanism Regulates CHIP E3 Ligase Activity

**DOI:** 10.1101/2021.08.20.457118

**Authors:** Aniruddha Das, Pankaj Thapa, Ulises Santiago, Nilesh Shanmugam, Katarzyna Banasiak, Katarzyna Dabrowska, Hendrik Nolte, Natalia A. Szulc, Rose M. Gathungu, Dominik Cysewski, Marcus Krüger, Michal Dadlez, Marcin Nowotny, Carlos J. Camacho, Thorsten Hoppe, Wojciech Pokrzywa

## Abstract

The E3 ubiquitin ligases CHIP/CHN-1 and UFD-2 team up to accelerate ubiquitin chain formation. However, it remained largely unclear how the high processivity of this E3 set is achieved. Here we studied the molecular mechanism and function of the CHN-1/UFD-2 complex in *Caenorhabditis elegans*. Our data show that UFD-2 binding promotes the cooperation between CHN-1 and ubiquitin-conjugating E2 enzymes by stabilizing the CHN-1 U-box dimer. The HSP-1 chaperone outcompetes UFD-2 for CHN-1 binding and promotes the auto-inhibited CHN-1 state by acting on the conserved position of the U-box domain. The interaction with UFD-2 enables CHN-1 to efficiently ubiquitinate S-Adenosylhomocysteinase (AHCY-1), an enzyme crucial for lipid metabolism. Our results define the molecular mechanism underlying the synergistic cooperation of CHN-1 and UFD-2 in substrate ubiquitylation.

**HIGHLIGHTS:**

- E3 ligase UFD-2 stimulates ubiquitylation activity of CHIP/CHN-1
- UFD-2 binding promotes dimerization of CHIP/CHN-1 U-box domains and utilization of E2 enzymes
- HSP70/HSP-1 by latching the U-box and TPR domains stabilizes the autoinhibitory state of CHIP/CHN-1, limiting interactions with E2s and UFD-2
- Assembly with UFD-2 enables CHIP/CHN-1 to regulate lipid metabolism by ubiquitylation of S-Adenosylhomocysteinase

## INTRODUCTION

The ubiquitin-proteasome system (UPS) includes a well-studied enzymatic cascade that transfers the small protein ubiquitin (Ub) onto a protein substrate (Kerscher et al., 2006). The last step in the UPS enzymatic cascade is mediated by ubiquitin-ligases (E3s), the largest and most diverse group of proteins within the UPS responsible for substrate selection and specificity (Komander, 2009; Buetow & Huang, 2016). In some instances, other proteins, i.e., ubiquitin chain elongation factors, or E4s, can be required to achieve efficient polyubiquitylation (poly-Ub) of model substrates. The first described E4 was yeast Ufd2p (Richly et al., 2005), a U-box domain-containing protein that engages Ub via its N-terminal region to assist with Ub chain elongation on pre-ubiquitylated substrates (Koegl et al., 1999; Hatakeyama et al., 2001; Buetow & Huang, 2016).

CHIP (C-terminus of Hsc70 interacting protein), initially identified as a tetratricopeptide repeat (TPR) protein that interacts with heat shock proteins (Ballinger et al., 1999), is a U-box E3 ubiquitin ligase that mediates ubiquitylation of chaperone client proteins, resulting in their degradation (Murata et al., 2001). Early *Caenorhabditis elegans* studies showed that UFD-2 interacts directly with CHN-1 (the nematode homolog of mammalian CHIP) to form an E3/E4 complex that can efficiently oligo-ubiquitylate the myosin chaperone UNC-45 (Hoppe et al., 2004). By contrast to the model proposed based on these early findings, more recent studies have revealed that UFD-2 acts as a true E3 ligase that poly-ubiquitylates UNC-45 independent of CHN-1, suggesting that both UFD-2 and CHN-1 act as E3s in the same or overlapping substrate space (Hellerschmied et al., 2018). A recent study aimed at identifying substrates of human CHIP and the human UFD-2 ortholog UBE4B supports the possibility of shared substrate scope (Bhuripanyo et al., 2018). However, despite the vital role of CHN-1/CHIP in protein quality control networks, little is known about its interactions with E3s and the regulation of its activity. To address these questions, we combined *in vitro* and *in vivo* assays with computational approaches and lipidomic and proteomic studies in *C. elegans* and uncovered the mechanism that controls CHN-1 activity. Our results indicate that UFD-2 interacts with the TPR domain of CHN-1 to boost CHN-1 processivity. This binding stabilizes the open conformation of the CHN-1 dimer enabling the U-box dimer to discharge more Ub-conjugating enzymes (E2) in a single ubiquitylation cycle. We also demonstrated that the heat shock protein HSP-1 interacts with the TPR and U-box domain of CHN-1 to stabilize the closed/auto-inhibitory state of CHN-1 dimer and limits its interaction with E2s and UFD-2. Furthermore, we identified potential substrates for the CHN-1/UFD-2 pair, including *S*-adenosylhomocysteinase (AHCY-1), a metabolic enzyme not known to be a client of known heat shock chaperones. Collectively, our results indicate an interplay between chaperones and UFD-2 in modulating CHIP activity. This processivity switching behavior of CHN-1 has important implications for its roles in regulating proteostasis, metabolism, and potentially other cellular processes.

## RESULTS

### UFD-2 promotes CHN-1 processivity and cooperation with E2s

Binding between CHN-1 and UFD-2 was previously demonstrated via yeast two-hybrid and *in vitro* pull-down assays (Hoppe et al., 2004). Beyond the physical interaction, the molecular regulation of Ub chain elongation by the CHN-1/UFD-2 complex has not been explored in detail. We performed an *in vitro* analysis of the activity of both E3s individually and in pairs. First, we chose E2s with which CHN-1 and UFD-2 cooperate in the auto-ubiquitylation (auto-Ub) reaction. Mammalian CHIPs can functionally interact with various E2s, particularly members of the UBCH5/UBE2D family (UBCH5a/UBE2D1, -b/2 and -c/3) (Jiang et al., 2001; Soss et al., 2011). Similarly, CHN-1 cooperates with UBE2D2 to mono-ubiquitylate (mono-Ub) *C. elegans* DAF-2 - insulin/insulin-like growth factor 1 (IGF-1) receptor *in vitro* (Tawo et al., 2017). To study the activity of CHN-1 in detail, we compared its ability to self-ubiquitylate in the presence of each of the UBE2D family proteins separately. We observed that CHN-1 cooperated effectively with UBE2D1. However, when we performed an auto-Ub reaction with both E3s, we observed a significant increase in CHN-1 poly-Ub activity, even when the E2 used in the reaction was UBE2D3, which is not efficiently used by CHN-1 alone (Figure 1A). Furthermore, the presence of UFD-2 also increases CHN-1 activity with the UBE2N/UBE2V1 E2 complex (Fig. S1B), which catalyzes the formation of free Ub chains that are then transferred to substrate proteins (Soss et al., 2011). We also concluded that the induction of E3 ligase activity is unidirectional since we did not detect any significant changes in the auto-Ub of UFD-2 under the same conditions (Fig. S1A). To gain insight into CHN-1processivity, we performed a time-dependent auto-Ub experiment. We observed no notable changes in the amount of ubiquitylated CHN-1 over time (from 60–180 min); however, the presence of UFD-2 strongly increased both mono- and poly-Ub of CHN-1 at the earliest time point (60 min) (Fig. 1B). Using the inactive CHN-1^H218Q^ mutant (Tawo et al., 2017), we observed that auto-Ub CHN-1 is not the result of modification by UFD-2 (Fig. S1C). Additionally, by deleting the TPR domain of CHN-1(Δ110aa), we confirmed its involvement in UFD-2 binding (Hoppe et al., 2004), as we could not observe modulation of the activity of this CHN-1 mutant by UFD-2 (Fig. S1D).

**Figure 1:**
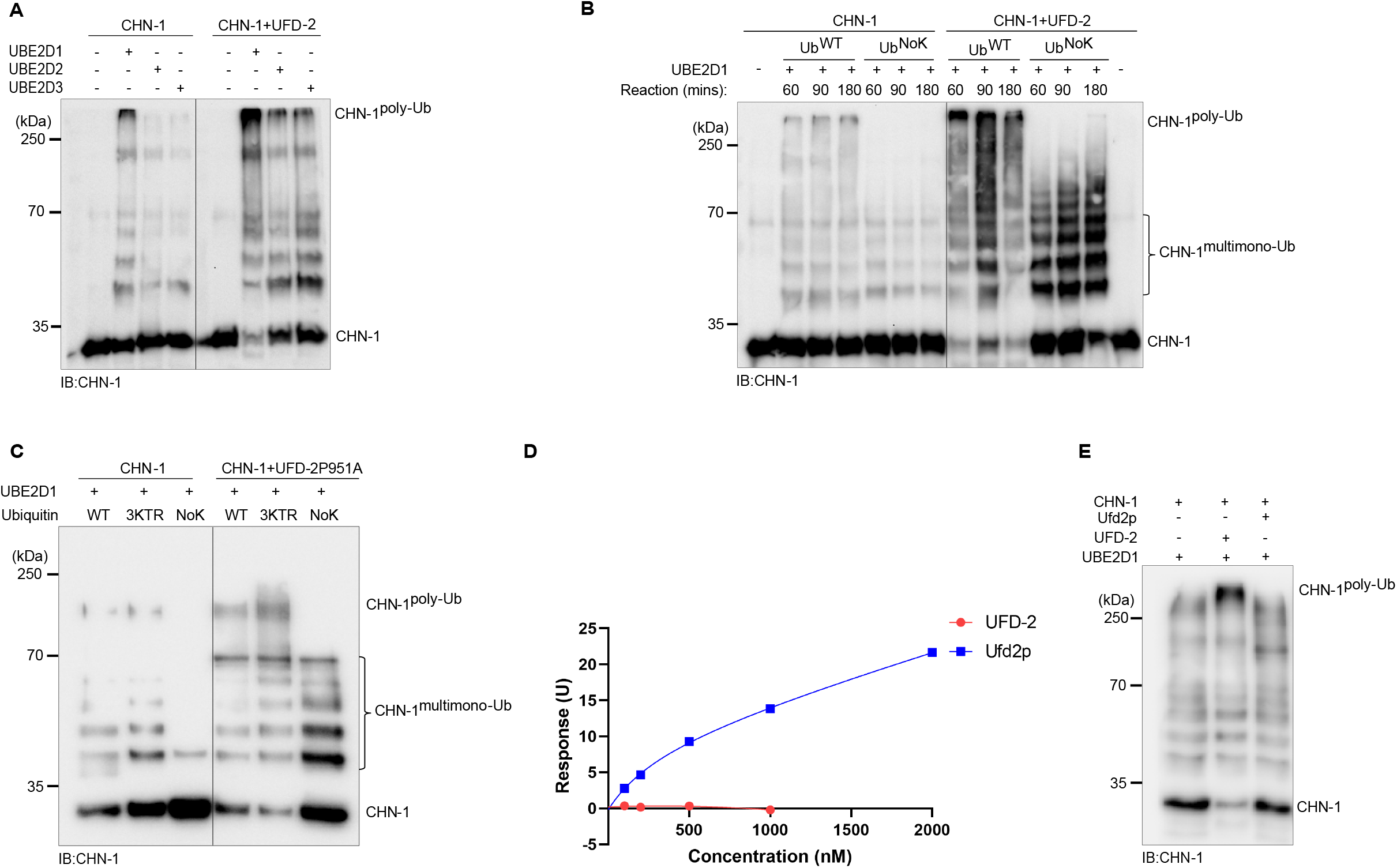
UFD-2 activates CHN-1. A) Auto-Ub of recombinant CHN-1 and UFD-2 was carried out using the E2s UBE2D1, UBE2D2, and UBE2D3. CHN-1 ubiquitylation was assessed via western blotting using CHN-1-specific antibodies. B) Time-dependent (60, 90, 180 min) auto-Ub of CHN-1 was performed as indicated using wild-type ubiquitin (Ub^WT^) or a lysine-free variant (Ub^NoK^). Protein samples were resolved via SDS-PAGE and immunoblotted with anti-CHN-1 antibodies. C) Auto-Ub was performed as indicated using recombinant CHN-1 and UFD-2^P951A^, UBE2D1 E2, Ub^WT^, Ub^NoK^ or Ub with substitutions of lysines 29, 48, 63 to arginines (Ub^3KTR^). Protein samples were resolved via SDS-PAGE and immunoblotted with anti-CHN-1 antibodies. D) Surface plasmon resonance (SPR) sensorgrams of the interaction between linear di-Ub (M1-linear from UbiQ) and *C. elegans* UFD-2 (red) or *S. cerevisiae* Ufd2p (blue). Y-axis: response unit (RU) value. X-axis: molar concentration of linear di-Ub. E) *In vitro* auto-ubiquitylation of CHN-1 in the presence of recombinant *C. elegans* UFD-2 and *S. cerevisiae* Ufd-2p, and UBE2D1 E2. Protein samples were resolved via SDS-PAGE and immunoblotted with anti-CHN-1 antibodies. Immunoblots representative of n = 3 experiments are shown.

Next, we wanted to verify whether UFD-2 can regulate the poly-Ub processivity of CHN-1 independent of its E3 activity. To examine this mechanism, we used an inactive, recombinant UFD-2 mutant with a P951A substitution (Ackermann et al., 2016). We found that UFD-2^P951A^ inactivity is due to its inability to bind an E2 enzyme (Fig. S1E). Finally, we performed a CHN-1 ubiquitylation reaction in the presence of UFD-2^P951A^. We detected substantial enhancement in both the mono- (using lysine-less Ub (UbK0)) and poly-Ub activity of CHN-1, regardless of the type of Ub chain (wild-type Ub or variant with substitutions of lysines 29, 48, 63 to arginines (UbKTR) (Fig. 1C). To rule out the possibility that UFD-2^P951A^ retained activity, we also used a UFD-2 variant (1–910 aa) lacking the entire U-box domain (909–984 aa). We confirmed that this UFD-2 deletion mutant could stimulate CHN-1 activity, indicating that interaction with some motif in UFD-2 alone is sufficient to activate CHN-1 (Fig. S1F).

Budding yeast Ufd2p can operate as a Ub chain elongation factor by interacting directly with Ub through its N-terminal region (Liu et al., 2017). Although higher eukaryotes have an ortholog of yeast Ufd2p, the Ub-interacting motif has little sequence homology (Hänzelmann et al., 2010; Liu et al., 2017) suggesting that the function of UFD-2 as an E4 is not evolutionarily conserved. To investigate whether the increased activity of the CHN-1/UFD-2 complex might stem from the elongation function of UFD-2, we tested whether UFD-2 retained the ability to interact with Ub using surface plasmon resonance (SPR) experiments. By contrast to Ufd2p, full-length UFD-2 did not bind linear Ub chains (Fig. 1D). This suggests that during evolution, UFD-2 lost its ability to elongate Ub chains directly. Unlike yeast Ufd2p, and perhaps to compensate for Ub binding loss, UFD-2 can induce processivity of its partner CHN-1 (Fig. 1E).

### UFD-2 induces structural gain of function in CHN-1

To gain mechanistic insight into the role of UFD-2 binding to CHN-1, we performed hydrogen-deuterium exchange mass spectrometry (HDX-MS) of the dimerization process of both CHN-1 alone and CHN-1 in the presence of UFD-2 (Fig. 2A and S2A). Available crystal structures of CHIP homologs support our HDX-MS analysis both without a chaperone (Nikolay et al., 2004) and with HSP90 (Zhang et al., 2005). In the absence of a TPR binding chaperone, only the dimer domains are revealed by the crystal structure, with no resolution of either the turn in the coil-coil domain or the TPR domain. Of note, the TPR by itself has only been resolved by NMR, whereas in the presence of an HSP substrate it stabilizes into its crystal form (Zhang et al., 2005). Furthermore, mouse CHIP shows that in one of its monomers, bound TPR is further stabilized against the long helix of its coil-coil domain. We have noted that this interaction is much weaker in CHN-1 (Thorsten Hoppe - personal communication), suggesting a more dynamic interaction in worms. Figure 2A and S2A depicts these states, leading to the following interpretation of our HDX-MS data, which detects at least three dynamical events at 10 s and 60 s. Namely, (a) the turn in the coil-coil motif (aa 146-152) is stabilized early on upon dimerization of the coil-coil domains; (b) the TPR domain is stabilized upon recognition by UFD-2, leaving the distal helices aa 21-40 and 92-112 exposed to solvent. At later times the stable TPR stabilizes against the long helix of the coil-coil domain; and, (c) the U-box domain (aa 21-40; 92-112) transitions from a weak interaction with its coil-coil domain to a stable dimer at longer time scales. As shown in Fig. 2A, we argue that contrary to CHIP (Ye et al., 2017; Zhang et al., 2005), CHN-1 folds into a symmetric structure as previously we have indicated that CHN-1 has critical residues that should prevent an asymmetric fold (Thorsten Hoppe - personal communication). Thus, while it has been shown that HSP90 negatively regulates CHIP activity (Narayan et al., 2015), presumably by blocking the E2 binding site of one of the protomers (Zhang et al., 2005), our findings of UFD-2 promoting CHN-1 processivity are consistent with a symmetric CHN-1 that upon binding UFD-2 stabilizes the U-box dimer with two E2 sites available for binding.

**Figure 2:**
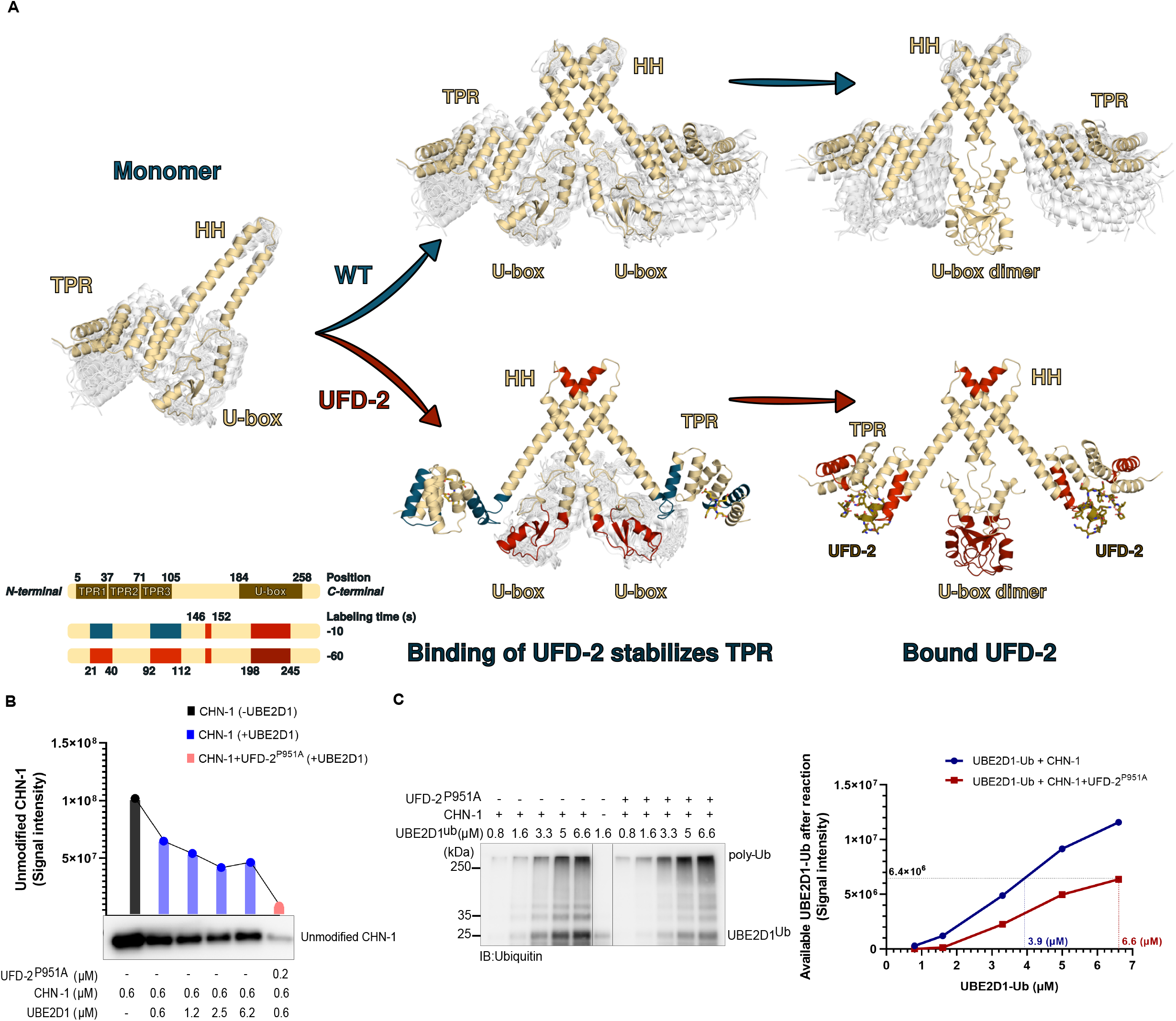
UFD-2 stabilizes CHN-1 U-box dimer. A) HDX-MS was used to analyze changes in the structural dynamics of residues within CHN-1 when in complex with UFD-2. The diagram model represents regions of retarded (red) and enhanced (blue) exchange in CHN-1 (upper panel). Schematics showing the domain organization of CHN-1 and the rate of deuterium exchange (colored box: blue, light red, medium red, dark red) in the different domains upon interaction with UFD-2 (lower panel). B) Auto-Ub of CHN-1 was performed as indicated using increasing amounts (0.6, 1.2, 2.5, 6.2 μM) of UBE2D1 or 0.6 μM UBE2D1 after complexing CHN-1 with 0.2 μM of recombinant UFD-2^P951A^. Protein samples were resolved via SDS-PAGE and immunoblotted with anti-CHN-1 antibodies. Quantification of the change in unmodified CHN-1 levels. Y-axis against the intensity of the unmodified band in each lane. Analysis performed using GraphPad Prism. C) E2 discharging assay of Ub-charged UBE2D1 in the presence of CHN-1/UFD-2^P951A^. The reaction was stopped after the indicated time via heat inactivation in native conditions. Protein samples were resolved via SDS-PAGE and immunoblotted with anti-Ub antibodies. (left panel). Quantification of available charged UBE2D1. Y-axis against the intensity of the UBE2D1-Ub signal from each and X-axis plotted against the µM concentration of UBE2D1-Ub (right panel). Immunoblots representative of n = 3 experiments are shown.

We conducted a ubiquitylation assay with increasing molar concentrations of UBE2D1 (0.6–6.2 µM) to confirm this model. We observed that at a constant Ub concentration, increasing E2 concentration led to an increase in CHN-1 activity. However, even at the highest E2 concentration (6.2 µM), CHN-1 processivity did not reach the same level as in the presence of inactive UFD-2^P951A^ and at an approximately 10-fold lower E2 concentration (0.6 µM) (Fig. 2B). Thus, the increased CHN-1 activity in the CHN-1/UFD-2 complex was not due to the increased local E2 concentration but rather to the enhanced processivity of the E2 enzyme bound to the CHN-1 U-box. To verify this hypothesis, we performed an E2-discharging assay in the presence of CHN-1 alone or after mixing with UFD-2^P951A^ to track the use of charged-E2 by CHN-1 only. We observed that in the presence of UFD-2^P951A^, CHN-1 could discharge almost twice as much of UBE2D1-Ub (approximately 6.6 µM) compared to CHN-1 alone (approximately 3.9 µM), as indicated by the accumulation of unused UBE2D1-Ub (Fig. 2C). Next, we performed another E2-discharging assay over time (0–30 min) to verify whether the increased utilization of charged E2 by the CHN-1/UFD-2^P951A^ pair was due to altered E2-E3 ubiquitin transfer dynamics. We noted that within the initial 5 minutes, the system achieved the maximum usage of charged E2, and no significant change in the level of available UBE2D1-Ub was observed over time (Fig S2B). Summarizing, our data indicate that UFD-2 acts as a preconditioner for the conformational flexibility of CHN-1, promoting dimerization of the U-box domains and thereby enabling their full functionality.

### HSP-1 and UFD-2 modulate CHN-1 processivity by stabilizing its inactive and active conformations, respectively

The three TPR domains in CHIP act as a binding platform for C-terminal peptides in the Hsp70 and Hsp90 chaperones, containing a conserved EEVD motif (Zhang et al., 2005; Paul & Ghosh, 2014; Zhang et al., 2015). Since CHN-1 also binds UFD-2 via the TPR domain, we investigated whether HSP-1 (the nematode Hsp70 orthologue) or DAF-21 (the nematode Hsp90 orthologue) could interfere with the activity of the CHN-1/UFD-2 complex. We first examined protein-protein interactions between CHN-1 and UFD-2, HSP-1, or DAF-21 using enzyme-linked immunosorbent assays (ELISAs). CHN-1 showed a higher affinity for HSP-1 and DAF-21 compared to UFD-2 (Fig. S3A). To verify the influence of HSP-1 and DAF-21 on the activity of the CHN-1/UFD-2 pair, we performed auto-Ub reactions in the presence of the chaperones. HSP-1 significantly reduced the auto-Ub activity of CHN-1 and blocked the stimulatory capacity of UFD-2 in this process (Fig. 3A). Negative regulation of CHIP auto-Ub by HSP70 has been previously reported, but the molecular basis is unclear (Narayan et al., 2015). Removal of the C-terminal EEVD motif deprived HSP-1 of its inhibitory effect. By contrast, DAF-21 did not affect the UFD-2-enhanced activity of CHN-1 (Fig. 3A).

**Figure 3:**
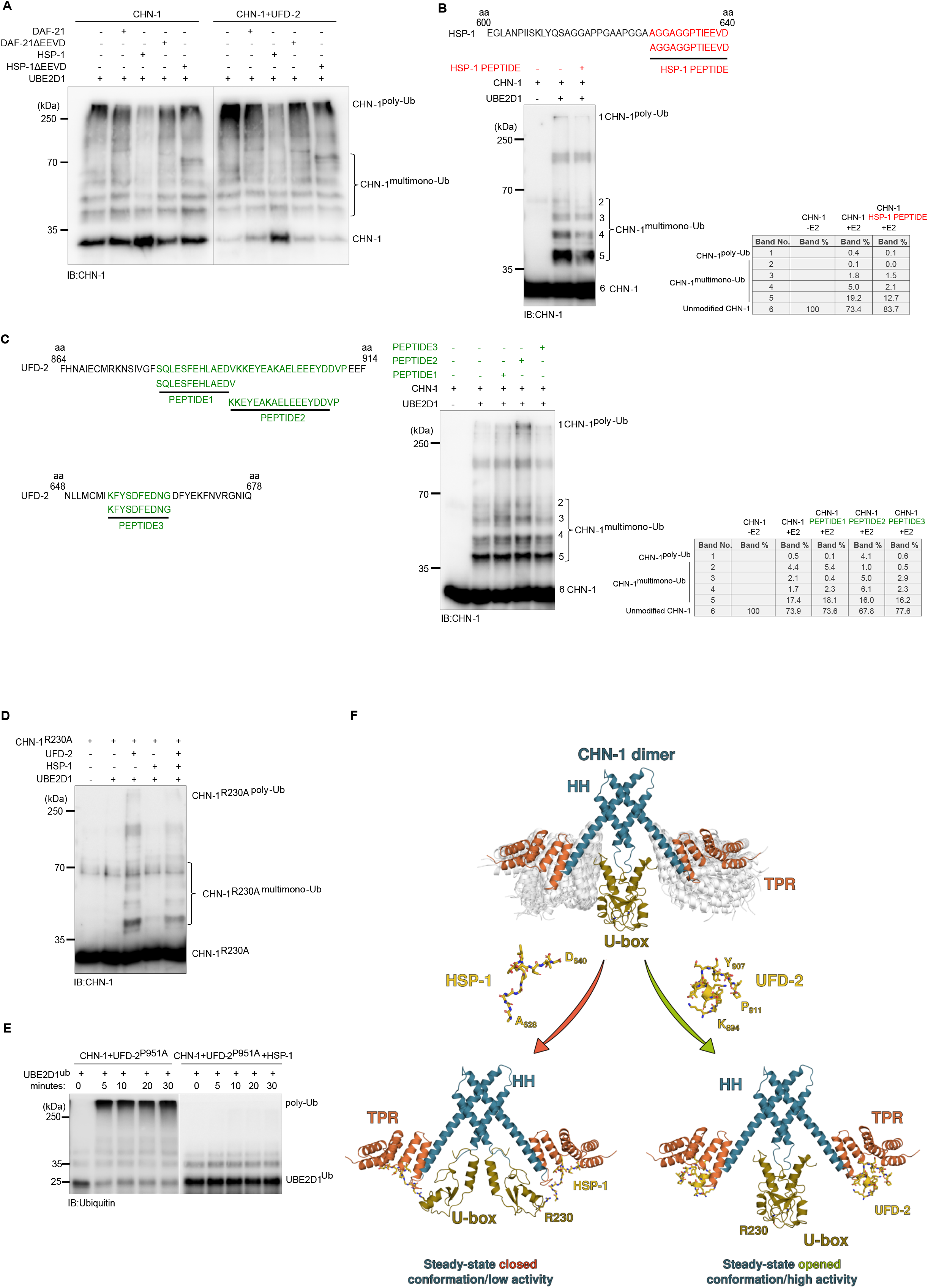
UFD-2 stabilizes an open/active, and HSP-1 stabilizes a closed/non-active CHN-1 conformation. A) *In vitro* auto-Ub was performed as indicated using recombinant CHN-1 complexed with UFD-2 in the presence of recombinant DAF-21, DAF-21ΔEEVD, HSP-1, or HSP-1ΔEEVD. Protein samples were resolved via SDS-PAGE and immunoblotted with anti-CHN-1 antibodies. B) On top schematics of the HSP-1 peptide sequence used in ubiquitylation reaction. Auto-Ub was performed as indicated using recombinant CHN-1 and HSP-1 derived peptide. Protein samples were resolved via SDS-PAGE and immunoblotted with anti-CHN-1 antibodies. Below, quantification of the changes (%) in (un)modified CHN-1 levels. Immunoblots representative of n = 3 experiments are shown. C) Schematics of the UFD-2 peptide sequences used in further ubiquitylation reactions (left panel). Auto-Ub was performed as indicated using recombinant CHN-1 and UFD-2 derived peptides. Protein samples were resolved via SDS-PAGE and immunoblotted with anti-CHN-1 antibodies (middle panel). Quantification of the changes (%) in (un)modified CHN-1 levels (right panel). D) Auto-Ub was performed as indicated using recombinant CHN-1^R230A^, UFD-2 and HSP-1. Protein samples were resolved via SDS-PAGE and immunoblotted with anti-CHN-1 antibodies. Immunoblots representative of n = 3 experiments are shown. E) E2 discharging assay of Ub-charged UBE2D1 in the presence of a ternary mixture of recombinant CHN-1/UFD-2^P951A^/HSP-1. The reaction was stopped after the indicated time via heat inactivation in native conditions. Protein samples were resolved via SDS-PAGE and immunoblotted with anti-Ub antibodies. F) Model of the UFD-2 activation and HSP-1 inhibition of CHN-1. Dimeric CHN-1 with TPR, U-box, and helix-turn-helix (HH) indicated by magenta, gold and cyan color, respectively. UFD-2 and HSP-1 peptides in yellow with indicated amino acid positions in the full-length proteins.

Next, we performed peptide mapping on peptide microarrays to pinpoint the interaction interface between the two ligases. For this, we used purified CHN-1 tagged with His::SUMO and His::SUMO alone (as control). These proteins were incubated on two UFD-2 peptide microarrays consisting of peptides of lengths 7 and 13 aa. This was followed by staining with secondary and control antibodies and reading using the LI-COR Odyssey Imaging System. Signal enrichment analysis suggested that the two consensus sequences EAKAELEEE and EEYDDVPE, were the predominant interactor motif. HSP70/90 uses a similar acidic C-terminal peptide with an EEVD sequence to bind to the TPR domain of target proteins (Scheufler et al., 2000; Gazda et al., 2013), and HSP-1 C-terminal EEVD peptide affected CHN-1 activity (Fig. 3B). Therefore, we examined whether the identified UFD-2 peptides could also regulate CHN-1. To this end, we performed CHN-1 auto-Ub reactions in the presence of the UFD-2-derived peptides identified in the peptide microarray data. We found that only the KKEYEAKAELEEEYDDVP peptide from UFD-2 significantly stimulated CHN-1 auto-Ub (Fig. 3C). An EEYD sequence is present in this peptide, suggesting that UFD-2 can utilize an EEVD-like motif for CHN-1 binding. Furthermore, multiple sequence alignment analysis revealed that in the EEYD motif of UFD-2, the amino acid Tyr (Y) is evolutionarily conserved among higher eukaryotes (Fig. S3B). To further define the functional role of the EEYD motif from UFD-2, we generated a chimeric recombinant HSP-1 protein carrying N-terminal EEYD instead of EEVD. Strikingly, we observed stimulation of CHN-1 auto-Ub when EEYD was introduced into HSP-1 – an opposite effect compared with that of wild-type HSP-1, which inhibited the reaction (Fig. S3C). Chimeric HSP-1^EEYD^ also exerted a slightly stimulatory effect on UBE2D1-Ub discharging by CHN-1 (Fig. S3D). This suggests that the CHN-1 activity switch can be regulated by its binding partners’ EEV(Y)D motifs.

To assess the contribution of the particular regions of the CHN-1 TPR domain, we generated its truncation variants. We showed that the first 87 amino acids (aa) (Δ87) are not responsible for the interaction with UFD-2 and HSP-1, and therefore are not involved in the modulation of CHN-1 processivity. In contrast, removing the subsequent eight residues (Δ95 variant) abrogated the CHN-1 poly-autoubiquitylation activity. Interestingly, the stimulating effect of UFD-2 was still observed, as evidenced by an increase in monoubiquitylated CHN-1^Δ95^ (Fig. S3E). CHN-1^Δ95^ has residues that may be involved in the interaction with UFD-2, including D110 and subsequent twists and helices (Fig. S3F). Indeed, the CHN-1^Δ110^ mutant, which lacks D110, does not show any gain of activity in the presence of UFD-2 (Fig S1D). It is known that a position homologous to D110 in mouse CHIP (D135) is involved in the binding of HSP’s, which suggests that this residue is also important for the interaction with UFD-2 EEYD peptide (Fig. S3F).

To understand why HSP-1 and UFD-2 peptides exhibit distinct effects on CHN-1 activity, we looked closely at the mechanism by which increased HSP90 and HSP70 concentration reduces CHIP activity (Narayan et al., 2015). As noted, HSP90 stabilizes an autoinhibit monomer in mouse CHIP (Zhang et al., 2005). Strikingly, this state entails a salt-bridge bridge between HSP90 D501 and CHIP R273, latching the U-box and TPR domains (Fig. S3G). This observation suggests that chaperone binding can directly restrain U-box from participating in Ub processivity. To show that a similar mechanism is at play in inhibiting ubiquitylation by HSP-1, we mutated R230 (homologous position to R273 in CHIP) to alanine to weaken the CHN-1 U-box interaction with the HSP-1 peptide and thus abrogate its inhibitory effect. Indeed, we observed significantly reduced inhibition of CHN-1R230A/UFD-2 complex by HSP-1 (Fig. 3D). Moreover, the addition of HSP-1 blocked the utilization of charged E2 by the CHN-1/UFD-2P951A complex (Fig. 3E). This agrees with the model indicating that by interacting with the TPR and U-box domain, HSP-1 stabilizes the autoinhibited state of CHN-1, affecting interaction with E2 enzymes (Fig. 3F). On the other hand, UFD-2 can avoid interacting with R230 by, for example, forming a helix that cannot extend toward the U-box, and induces uncorrelated mobility of the TPR domains with respect to the U-box domains, promoting the steady-state open conformation of CHN-1 (Fig. 3F).

### The CHN-1/UFD-2 pair regulates phosphatidylcholine synthesis via AHCY-1

We next wished to establish the functional consequences of CHN-1/UFD-2 cooperation *in vivo*. Based on our *in vitro* studies, we hypothesized that *in vivo* CHN-1, when functioning alone, would display insufficient poly-Ub activity and mainly mono-Ub substrates. Indeed, Tawo and colleagues showed that CHN-1/CHIP mono-Ub the DAF-2 insulin receptor (Tawo et al., 2017). We further assumed that the interaction with UFD-2 would trigger the poly-Ub activity of CHN-1, consequently leading to efficient degradation of its specific substrates. Thus, to understand the role of CHN-1 and UFD-2 *in vivo*, we decided to identify such substrates. We searched for proteins whose levels increase after deletion of CHN-1 (substrate ubiquitylation by CHN-1 would be affected directly) or UFD-2 (CHN-1 would not be stimulated to efficiently poly-Ub its substrates). To define the consequences of *chn-1* and *ufd-2* deletion on the *C. elegans* proteome and to detect proteins that accumulate in the deletion mutants in an unbiased way, we performed label-free mass spectrometry (LC-MS/MS)-based proteomics experiment. We analyzed *chn-1(by155), ufd-2(tm1380)*, and *chn-1(by155); ufd-2(tm1380)* double-mutant worms by single-shot LC-MS/MS gradients in 5 biological replicates. To obtain a view of the global structure of the data, we performed dimensional reduction using principal component analysis (PCA). We noticed that the proteomes of *chn-1(by155)* and *chn-1(by155); ufd-2(tm1380)* mutants clustered closer together than did those of *chn-1(by155)* and *ufd-2(tm1380)* (Fig. S4A). We hypothesized that potential substrates should accumulate in all mutants; therefore, we filtered the set of significantly altered proteins requiring a two-fold enrichment in all mutants versus the N2 control strain. We obtained 65 potential substrate candidates and visualized them via hierarchical clustering (Fig. S4B, C and Supp. Table 1). These potential substrates were enriched in metabolic processes, including lipid biosynthesis, as shown via Gene Ontology over-representation analysis (Fig. S4D), and among them, we identified the AHCY-1 enzyme (Fig. 4A and S4C). AHCY-1 catalyzes the reversible hydrolysis of *S*-adenosylhomocysteine (SAH) to homocysteine and adenosine (Palmer and Abeles, 1976; 1979) (Fig. 4B). Despite the fundamental role of AHCY-1 in metabolism, its regulatory mechanisms are still enigmatic. In a yeast two-hybrid screen using a *C. elegans* cDNA library, we identified AHCY-1 as the prominent interactor of CHN-1 (Fig. S4E). We confirmed the interaction between the two proteins in worms via co-immunoprecipitation (Fig. S4F). Next, we tested whether AHCY-1 is a CHN-1 substrate by performing *in vitro* ubiquitylation assays with recombinantly expressed proteins. We confirmed that recombinant AHCY-1 is a specific substrate of CHN-1 that UFD-2 does not ubiquitylate (Fig S4H). Furthermore, in the presence of UFD-2, CHN-1 poly-Ub AHCY-1 more effectively, and the level of this modification was reduced by HSP-1 (Fig. 4C, S4G). The cooperation between CHN-1 and UFD-2 is also consistent with the detection of a similar increase in the AHCY-1 level in *chn-1(by155), ufd-2(tm1380)*, and double mutant worms in our proteomic analysis (Fig. 4A). To further validate this observation, we monitored the endogenous level of AHCY-1 via western blotting of total lysates of wild-type worms, *chn-1(by155)* and *ufd-2(tm1380)* mutant worms, and worms overexpressing *chn-1* treated with the proteasome (MG132) and DUB (*N*-methylmaleimide, NEM) inhibitors. We did not observe any significant changes in the AHCY-1 level, which, according to our other observations, is a stable and abundant protein in *C. elegans*. However, immunoblotting analysis with anti-AHCY-1 antibodies detected higher molecular weight smeared bands, likely corresponding to polyubiquitinated AHCY-1 species. These AHCY-1 modifications were more abundant when *chn-1* was over-expressed and were reduced in *chn-1(by155)* and *ufd-2(tm1380)* mutant worms compared with the ACHY-1 status in wild-type animals (Fig. 4D). These data suggest that CHN-1 regulates AHCY-1 *via* an E3 activity triggered by UFD-2.

**Figure 4:**
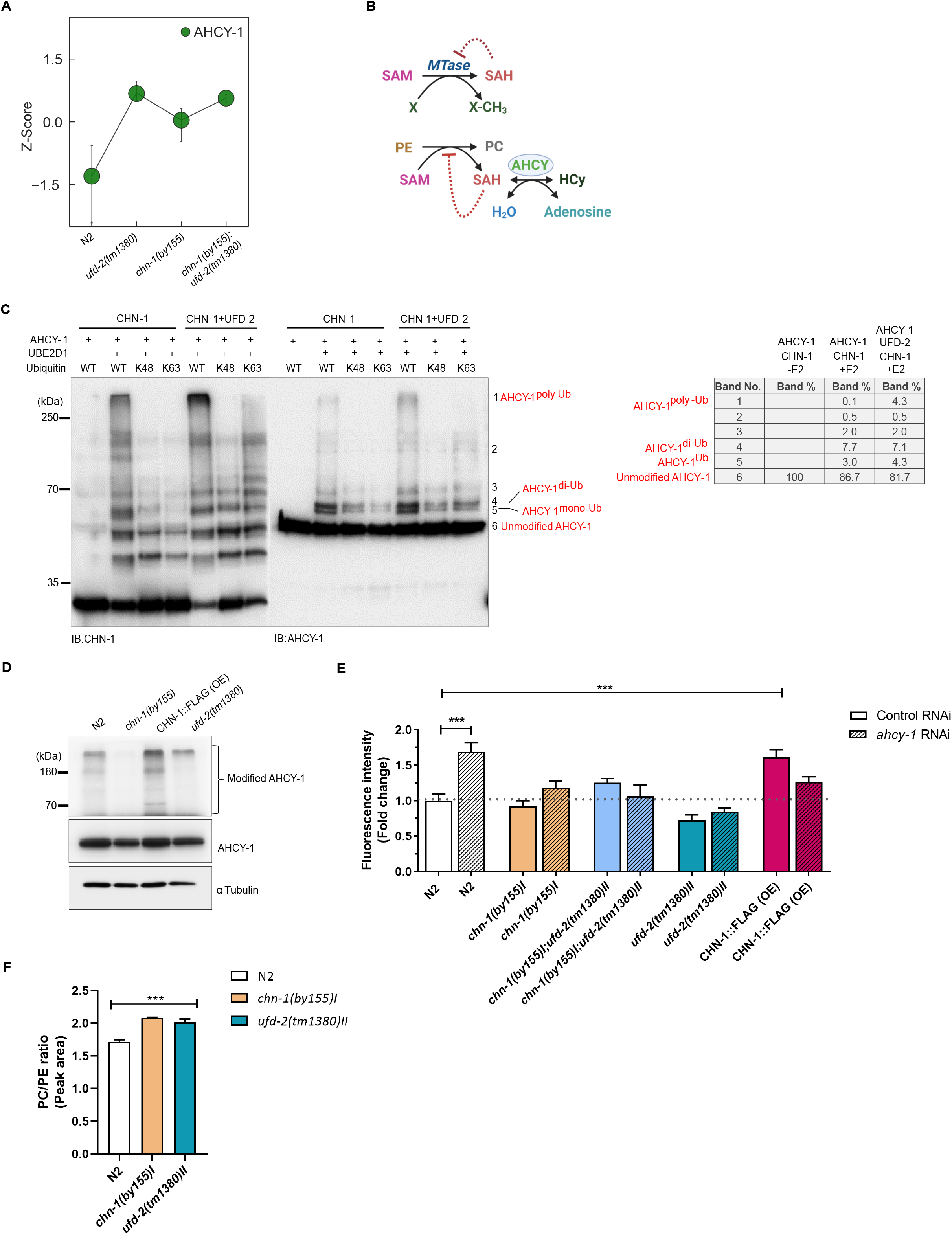
The CHN-1/UFD-2 pair regulates lipid metabolism via AHCY-1. A) Endogenous levels of AHCY-1 in N2 (wild-type), *chn-1(by155), ufd-2(tm1380)*, and *chn-1(by155); ufd-2(tm1380)* mutant worms reported as Z-scores from LC-MS/MS analysis. B) Schematic diagram representing the core function of AHCY. AHCY catalyzes the reversible hydrolysis of SAH (S-adenosylhomocysteine) to HCy (Homocysteine). Accumulation of SAH inhibits PC (Phosphatidylcholines) synthesis from PE (Phosphatidylethanolamines). C) Ubiquitylation of recombinant AHCY-1 was performed as indicated using recombinant CHN-1 and UFD-2, UBE2D1 E2, Ub^WT^, or Ub^K48^ and Ub^K63^ only Ub variants. Protein samples were resolved via SDS-PAGE and immunoblotted with anti-AHCY-1 antibodies. Bands labeled as unmodified AHCY-1, mono-Ub AHCY-1, di-Ub AHCY-1, poly-Ub AHCY-1 (left panel). Quantification of the changes (%) in (un)modified AHCY-1 levels (right panel). D) Endogenous levels of AHCY-1 in N2 (wild-type), *chn-1(by155)*, CHN-1::FLAG (OE), and *ufd-2(tm1380)* young adult worms treated with a proteasome inhibitor (MG132, 10μM) or DUB inhibitor (NEM, 100mM). Protein samples were resolved via SDS-PAGE and immunoblotted with anti-AHCY-1 antibodies. Tubulin served as a loading control. Immunoblots representative of n = 3 experiments are shown. E) Total lipid content in N2 (wild-type), *chn-1(by155), ufd-2(tm1380), chn-1(by155), ufd-2(tm1380)*, and CHN-1::FLAG (OE) young adult worms grown on control and *ahcy-1* RNAi feeding plates. Data are means ± SEM, p ≤ 0.001 (***). Higher fluorescence intensity indicates increased lipid levels. F) Ratio of phosphatidylcholine (PC) to phosphatidylethanolamine (PE) in N2 (wild-type), *chn-1(by155)*, and *ufd-2(tm1380)* young adult worms.

Elevated homocysteine levels are linked to the deregulation of lipid metabolism and increased fat accumulation, apparent after RNA interference (RNAi) depletion of AHCY-1 in worms (Vrablik et al., 2015; Visram et al., 2018). Using the lipophilic fluorophore RediStain WormDye Lipid Green to stain and quantify the fat content of *C. elegans*, we confirmed that AHCY-1 depletion increases the abundance of lipids in wild-type worms by almost 60%. Overexpression of *chn-1* caused an increase in total lipid content to the similar level detected in *ahcy-1* RNAi-treated worms, and this effect was not further enhanced by AHCY-1 depletion (Fig. 4E). Interestingly, mutations in *chn-1* and *ufd-2* cause a reduction in overall lipid levels and uncouple the stimulation of lipid biogenesis induced by *ahcy-1* RNAi (Fig. 4E). Synthesis of phosphatidylcholine (PC) from phosphatidylethanolamine (PE) via the *de novo* phospholipid methylation pathway requires a significant amount of SAM and is particularly sensitive to SAH levels (Tehlivets, 2011). Consistent with our assumption that deletion of either *chn-1* or *ufd-2* would positively affect AHCY-1 stability, leading to intensification of SAM-dependent methylation and PE to PC conversion, we noted that the ratio of PC to PE increased in *chn-1(by155)* and *ufd-2(tm1380)* worms (Fig. 4F). In conclusion, our data suggest a functional role for the CHN-1/UFD-2 complex in AHCY-1-dependent lipid metabolism regulation.

## DISCUSSION

The different conformations achieved by dynamic and flexible motifs are important for the functionality of various E3 ligases (Faull et al., 2019; Kamadurai et al., 2013; Liu and Nussinov, 2011; Narayan et al., 2015). The crystal structure of murine CHIP E3 bound to an HSP-90 decapeptide containing an EEVD motif revealed an asymmetric dimerization in which the two CHIP protomers adopt different conformations. In this “closed” state, only one of the U-box domains in the dimer is accessible for E2 binding, and the other is blocked by the TPR domain (Zhang et al., 2005). In agreement with a computational model of human CHIP (Ye et al., 2017), our homology modeling of the CHN-1 dimer suggested that it can take the form of both a metastable symmetric dimer in which both U-box domains can simultaneously bind E2 ubiquitin-conjugating enzymes and asymmetric dimer with low ubiquitylation activity. We showed that the interaction of E3 UFD-2 with the CHN-1 TPR domain reduces its dynamics and thus its blocking of U-box domains. In this steady-state open conformation, CHN-1 achieved high poly-Ub activity due to the full functionality of the U-box dimer. Consistently, in the E2 discharging assay, we observed a two-fold increase in the utilization of charged E2 by the CHN-1/UFD-2^P951A^ complex compared with that of CHN-1 alone. Here, UFD-2 acts as a pre-conditioning factor to influence the conformational flexibility of CHN-1, allowing higher processivity at the initial phase; therefore, we did not observe any further change in our E2 discharging reaction kinetics with increasing time. Using various ubiquitin variants, we showed that not only poly-Ub but also CHN-1 mono-Ub, which is the rate-limiting step of ubiquitylation, is also enhanced upon UFD-2 binding. We also found that UFD-2 activity is unaffected in the complex and that the two ligases are not substrates for each other.

The N-terminal TPR domain of CHIP has been shown to interact specifically with the C-terminal EEVD motif of HSP70 and HSP90 (Zhang et al., 2005; Xu et al., 2006; Graf et al., 2010). We discovered that UFD-2 uses a slightly modified motif - EEYD to engage the CHN-1 TPR domain. Furthermore, we demonstrated that only the presence of a UFD-2 peptide containing the EEYD sequence was sufficient to promote CHN-1 activity. In contrast, the *C. elegans* HSP70 homolog, HSP-1, negatively regulates CHN-1 and CHN-1/UFD-2 complex activity by promoting its auto-inhibited (closed) CHN-1 state. GHFDPVTR sequence in the U-box domain is evolutionarily conserved in CHIP from different species, but its role was not previously known. Here we showed that CHN-1 activity is negatively regulated by the interaction between positions associated with the EEVD motif of HSP-1 and the conserved R230 position in the GHFDPVTR sequence. Through direct interactions with the TPR and U-box domain of CHN-1, HSP-1 brings both regions to proximity impairing the U-box dimer. This depends only on the local interaction of the C-terminus of HSP-1 with the U-box, not on the steric hindrance of E2 access to U-box domains by the whole chaperone. In co-crystal with CHIP, HSP90 also forms hydrogen bonds (H-bonds) between T and S in its C-terminal peptide (TSRMEEVD) and TPR of CHIP (Zhang et al., 2005). The existence of these H-bonds between the HSP-1 peptide (GPTIEEVD) and CHN-1 is not apparent. However, the C-terminal sequence of HSP-1 is rich in glycines that may tailor the binding more efficiently by forming H-bonds with the CHN-1 backbone, possibly leading to a very close interaction. Accordingly, the C-terminal HSP70 peptide blocks CHIP activity markedly greater than the HSP90 peptide, which binds to the CHIP TPR domain weaker than the HSP70 peptide (Narayan et al., 2015). We also observed that worm DAF-21/HSP90 has a lower affinity for CHN-1 and does not affect CHN-1 activity, unlike HSP-1. Thus, the mechanism in which the degree of interaction depends on the C-terminal sequence of the chaperones is conserved and correlates with the stabilization of the autoinhibited CHN-1/CHIP dimer. Presumably, high CHN-1 processivity is undesirable for HSPs as it could lead to an imbalance between chaperone-mediated folding/maturation and degradation, inducing the latter. We cannot exclude the influence of posttranslational factors or the cellular environment on the level of regulation of CHN-1 activity by HSP-1 and UFD-2.

Previous research has established that CHIP participates in protein quality control by routing a wide range of chaperone substrates for degradation (Joshi et al., 2016). Our observations suggest an alternative, non-quality control role for the CHN-1/UFD-2 complex. We identified AHCY-1 as a novel substrate of the CHN-1/UFD-2 complex. AHCY-1 is the only eukaryotic enzyme capable of hydrolyzing SAH, which is essential for SAM-dependent methylation (Cantoni, 1975). Recent findings support the importance of CHIP in regulating the methylation status of the cellular proteome by mediating proteasomal turnover of the SAM-dependent methyltransferases PRMT1, PRMT5, and EZH2 (Zhang et al., 2016; Bhuripanyo et al., 2018). However, further studies are necessary to delineate the involvement of the CHN-1/UFD-2 complex in modulating the cellular methylation potential. In summary, our data provide mechanistic insights into the distinct regulation of CHN-1/CHIP activity by HSP70 and UFD-2 and the processivity of non-chaperone CHIP substrates.

## LIMITATIONS OF THE STUDY

Our study elucidates the possible mechanism of action of CHN-1 in the presence of various partner proteins based on the biochemical data and the available structure of CHIP. However, we were unable to provide detailed insights into the structure of CHN-1 in association with UFD-2 or HSPs as we were unable to obtain a co-crystal after various attempts. Furthermore, we could not isolate the essential amino acids of CHN-1 that are required for interaction with UFD-2 and HSP-1. Therefore, we cannot exclude the possibility that multiple CHN-1 motifs may be involved in the interactions. In addition to presenting a novel, non-quality-controlling role for the CHN-1/UFD-2 pair, we cannot comment on the physiological factors that regulate this E3 complex assembly.

## LEGENDS

**Supplementary Figure S1:** A) Auto-Ub of UFD-2 was performed as indicated using E2s UBE2N/Uev1a, UBE2D1, or UBE2D3, and Ub^WT^ or Ub^NoK^. Protein samples were resolved via SDS-PAGE and immunoblotted with anti-UFD-2 antibodies. B) Auto-Ub was performed as indicated using UBE2N/Uev1a and UBE2D1 E2s. Protein samples were resolved via SDS-PAGE and immunoblotted with anti-Ub antibodies. C) Ubiquitylation of recombinant CHN-1^H218Q^ mutant was performed as indicated. Protein samples were resolved via SDS-PAGE and immunoblotted with anti-CHN-1 antibodies. D) Auto-Ub of recombinant CHN-1^Δ110^ was performed as indicated using UBE2D1 E2. Protein samples were resolved via SDS-PAGE and immunoblotted with anti-CHN-1 antibodies. E) Co-immunoprecipitation of ubiquitin-charged GST-UBE2D1 from a mixture of recombinant GST-UBE2D1 and CHN-1, GST-UBE2D1 and UFD-2^P951A^, and the ternary mixture of GST-UBE2D1, CHN-1 and UFD-2^P951A^ using Dynabeads conjugated with anti-GST antibody. Protein samples were resolved via SDS-PAGE and immunoblotted with anti-GST, anti-UFD-2, and anti-CHN-1 antibodies. F) Auto-Ub was performed as indicated using recombinant UFD-2, UFD-2^P951A^, or UFD-2^ΔUbox^. Bands labeled as unmodified CHN-1, mono-Ub CHN-1, and poly-Ub CHN-1. Below, quantification of the changes (%) in (un)modified CHN-1 levels. Immunoblots representative of n = 3 experiments are shown.

**Supplementary Figure S2:** A) Differential Woods plots present the difference in fractional deuterium uptake between two biological states - CHN-1 and CHN-1 in the presence of wild-type UFD-2. The X-axis represents the position in sequence for a peptide (the x value indicates the peptide length). The Y-axis presents the difference in fractional deuterium uptake with the Y-error bar indicating the uncertainty of the measurement from three independent replicates of the experiment. Positive values indicate stabilization of the region upon complex formation. Dotted lines indicate the confidence limit at 95% calculated using the Houde test (Houde et al., 2011). The upper and lower panels show results after 10 and 60 seconds of H/D exchange, respectively. B) E2 discharging assay of Ub-charged UBE2D1 by the recombinant CHN-1/UFD-2^P951A^. The reaction was stopped after the indicated time via heat inactivation in native conditions. Protein samples were resolved via SDS-PAGE and immunoblotted with anti-Ub antibodies.

**Supplementary Figure S3:** A) Titration ELISA assay to determine the dissociation constant (K_D_) between DAF-21, HSP-1, UFD-2, and CHN-1. Y-axis: CHN-1 concentration (µM). X-axis: absorbance (OD) at 450 nm. Below, a table showing the K_D_ value (nM) of the corresponding protein with recombinant CHN-1. B) Multiple sequence alignment (MSA) of UFD-2 orthologs from different species. Orthologous sequences (from Orthologous Group ID ENOG5038DSP) of selected species were obtained from the eggNOG5 database (Huerta-Cepas et al., 2019) and aligned using the T-Coffee web server with default parameters (Di Tommaso et al., 2011; Notredame et al., 2000). Vertebrates possess two UFD-2 orthologs, which have been independently annotated. MSA was visualized in Jalview Desktop software (Waterhouse et al., 2009) with residues colored according to their physicochemical properties; conserved tyrosine (Y) residues and the EEYD motif in *C. elegans* are highlighted in white frames. C) Auto-Ub was performed as indicated using recombinant CHN-1 complexed with HSP-1 and HSP-1^EEYD^. Protein samples were resolved via SDS-PAGE and immunoblotted with anti-CHN-1 antibodies. Immunoblots representative of n = 3 experiments are shown. D) E2 discharging assay of Ub-charged UBE2D1 in the presence of CHN-1 or CHN-1/HSP-1^EEYD^. The reaction was stopped after the indicated time via heat inactivation in native conditions. Protein samples were resolved via SDS-PAGE and immunoblotted with anti-Ub antibodies. Immunoblots representative of n = 3 experiments are shown. E) *In vitro* autoubiquitylation was performed as indicated using recombinant CHN-1^Δ87^ or CHN-1^Δ95^ truncation mutants in the presence of UFD-2, DAF-21, DAF-21ΔEEVD, HSP-1, and HSP-1ΔEEVD. Samples were analyzed by SDS-PAGE and immunoblotted with anti-CHN-1 antibodies. Immunoblots representative of n = 3 experiments are shown. F) Model of the CHN-1 TPR domain docked with UFD-2 EEYD peptide with 1-86 residues of CHN-1 colored in orange and 87-95 residues in magenta colors that sequester the EEYD motif away from R230 residue of CHN-1. G) Co-crystal structure of the mice CHIP TPR domain showing interaction with HSP90 EEVD peptide (2C2L) shows R273 (conserved in CHN-1 as R230) in proximity close enough to interact with D501 of HSP90.

**Supplementary Figure S4:** A) PCA analysis showing the first and second principal components of the significantly altered proteins (ANOVA FDR < 0.05) performed in the Perseus software. The percentage of explained variance is provided on the axis labels as a percentage. B) Schematic representation of the number of identified proteins in a single-shot analysis of LC-MS/MS gradients in 5 biological replicates that led to the identification of proteins with a significant change in abundance in *chn-1(by155), ufd-2(tm1380)*, and *chn-1(by155); ufd-2(tm1380)* worms (two-fold enrichment in all mutants versus N2 (wild-type) animals). C) Hierarchical clustering of the Z-Score of proteins whose levels increased in *chn-1(by155), ufd-2(tm1380)*, and *chn-1(by155); ufd-2(tm1380)* mutant worms (two-fold enrichment in all mutants versus N2 (wild-type) animals from LC-MS/MS experiment). D) Gene Ontology biological process terms found to be associated with *C. elegans* genes upregulated (minimum two-fold enrichment versus N2 (control), with FDR < 0.05 for ANOVA or pairwise t-test) in all mutants; all proteins detected in LC-MS/MS analysis constituted a reference set. Over-representation analysis was performed using the WebGestalt web server with default parameters (Liao et al., 2019). FDR was controlled to 0.25 using the Benjamini-Hochberg method for multiple testing. E) Yeast 2-hybrid prey fragment analysis. Schematic representations of the AHCY-1 fragments interacting with CHN-1. The coding sequence for CHN-1 was used as bait to screen a random-primed *C. elegans* mixed-stage cDNA library. The selected interaction domain (SID) is the amino acid sequence shared by all AHCY-1 fragments (prey) interacting with CHN-1. The confidence score of this binding (predicted biological score) is A (highest confidence). F) Co-immunoprecipitation of AHCY-1 and UFD-2 from young adult worms expressing CHN-1::FLAG using beads conjugated with anti-FLAG antibody. Protein samples were resolved via SDS-PAGE and immunoblotted with anti-AHCY-1, anti-FLAG, and anti-UFD-2 antibodies. (The red boxes mark the protein band). G) Ubiquitylation of recombinant AHCY-1 was performed as indicated using recombinant CHN-1, UFD-2, DAF-21, HSP-1 in the presence of UBE2D1. Protein samples were resolved via SDS-PAGE and immunoblotted with anti-AHCY-1 antibodies. H) Ubiquitylation of recombinant AHCY-1 was performed as indicated using recombinant UFD-2 and UBE2D1 E2. Protein samples were resolved via SDS-PAGE and immunoblotted with anti-AHCY-1 antibodies. Immunoblots representative of n = 3 experiments are shown.

## STAR★METHODS

### Lead Contact

Further information and requests for reagents may be directed to and will be fulfilled by Wojciech Pokrzywa.

### Materials Availability

Plasmids generated by the authors will be distributed upon request to other researchers.

### Data a Availability

The mass spectrometry proteomics data was deposited to the ProteomeXchange Consortium via the PRIDE partner repository with the dataset identifier PXD028023 (Perez-Riverol et al., 2019).

### Generation of recombinant proteins

All recombinant proteins were produced using a bacterial expression system. CHN-1 and the CHN-1 variants were expressed and purified from Rosetta™ 2 (DE3) cells. UFD-2, HSP-1, DAF-21, and their variants were expressed and purified from BL21 Star™ (DE3) cells. Truncations and point mutations in the protein constructs were introduced using the Q5 Site-Directed Mutagenesis Kit (NEB, Cat#E0552S). Protein expression was induced using 0.4 mM IPTG at 22 °C for 16 hr. Respective induced cell pellets were harvested via centrifugation at 4000 rpm for 20 min at 4 °C. Cells were lysed in a lysis buffer (20 mM HEPES pH 8, NaCl 300 mM, 2 mM BME, protease inhibitor, and DNase) by sonication. After sonication, the supernatant and pellet fractions were separated via high-speed centrifugation at 14000 rpm for 1 hr at 22 °C. Tagged proteins were purified from the soluble fraction of the cell lysates using appropriate Ni-NTA or GST Hi-trap columns or chitin beads (NEB, Cat#E6901S). After removing the affinity tags, affinity-purified protein fractions were subjected to gel filtration chromatography (Hiload 16/600 Superdex S200, GE Healthcare) to obtain more than 95% pure protein fractions for use in subsequent biophysical and biochemical experiments. For the *in vitro* ubiquitylation reactions, we first generated a pTYB21-UFD-2 expression vector and purified tagless UFD-2 fraction using the intein cleavage site as per the manufacturer protocol (NEB Cat#E6901S). The lysis buffer used for purifying this variant contained HEPES 20mM, TritonX 0.1%, 5% glycerol, 500 mM NaCl, pH 8.0. To generate tagless CHN-1 and His-tagged CHN-1, we affinity-purified the proteins using Ni-NTA columns and then collected the dimeric fraction using size-exclusion chromatography (SEC). Furthermore, SUMO and the His-tag were cleaved using SUMO protease treatment (16 hr) at 4 degrees, and untagged CHN-1 was purified via SEC.

### Peptide microarray for protein-peptide interaction studies

This assay was performed by PEPperPRINT GmbH (https://www.pepperprint.com/). The CHN-1 and UFD-2 sequences were elongated with neutral GSGSGSG linkers on the C- and N-termini to avoid truncated peptides. The elongated CHN-1 sequence was translated into 7, 10, and 13 amino acid peptides with peptide-peptide overlaps of 6, 9, and 12 amino acids. The elongated UFD-2 sequence was translated into 7 and 13 amino acid peptides with peptide-peptide overlaps of 6 and 12 amino acids. After peptide synthesis, all peptides were cyclized via a thioether linkage between a C-terminal cysteine and an appropriately modified N-terminus. The resulting conformational CHN-1 and UFD-2 peptide microarrays contained 813 and 1,986 different peptides printed in duplicate (1,626 and 3,972 peptide spots), respectively. The peptide array was framed by additional HA control peptides (YPYDVPDYAG, 68 spots for CHN-1 and 130 spots for UFD-2). Samples: His-tagged SUMO CHN-1, His-tagged SUMO and His-tagged UFD-2 proteins. Washing Buffer: TBS, pH 7.5 with 0.005% Tween 20; washing for 2 × 10 sec after each incubation step. Blocking Buffer: Rockland blocking buffer MB-070 (30 min before the first assay). Incubation Buffer: TBS, pH 8 with 10% Rockland blocking buffer MB-070, 10 mM HEPES, 150 mM NaCl and 0.005% Tween 20. Assay Conditions: Protein concentrations of 10 µg/mL and 100 µg/mL in incubation buffer; incubation for 16 h at 4 °C and shaking at 140 rpm. Secondary Antibody: Mouse anti-6x-His Epitope Tag DyLight680 (1.0 µg/mL); 45 min staining in incubation buffer at RT. Control Antibody: Mouse monoclonal anti-HA (12CA5) DyLight800 (0.5 µg/mL); 45 min staining in incubation buffer at RT. Scanner: LI-COR Odyssey Imaging System; scanning offset 0.65 mm, resolution 21 µm, scanning intensities of 7/7 (red = 700 nm/green = 800 nm). Microarray Data: Microarray Data Sumo Protein (PEP20205011547).xlsx, Microarray Data Sumo CHN-1 Protein (PEP20205011547).xlsx, Microarray Data UFD-2 Protein (PEP20205011547).xlsx. Microarray Identifier: 002413_05 (CHN-1 microarray, four array copies for one-by-one assays) 002413_07 & 002413_08 (UFD-2 microarray, two array copies for one-by-one assays).

### Ubiquitylation assays

*In vitro* assays were performed according to an earlier protocol (Hellerschmied et al., 2018). The reactions were run at 30 °C for 90 minutes using 60 µM Ubiquitin (Boston Biochem, Ub^WT^ Cat#U-100H; Ub^NoK^, Cat#UM-NOK; Ub^3KTR^, Cat#UM-3KTR; Ub K48 only, Cat#UM-K480; Ub K63 only, Cat#UM-K630) in the presence of 100 nM E1 (UBE1, Boston Biochem, Cat#E-304), 0.6 µM E2 (Boston Biochem, UBE2D1, Cat#E2-616; UBE2D2, Cat#E2-622; UBE2D3, Cat#E2-627; UBE2N/Uev1a, Cat#E2-664), E3 ligase (1 µM CHN-1 and variants or 0.7 µM UFD-2 and variants), E3 ligase reaction buffer (Boston Biochem, Cat#B-71), and Energy Regeneration Solution (Boston Biochem, Cat#B-10). For performing the *in vitro* reaction in the presence of both the CHN-1 and UFD-2 or His-tagged UFD-2^P951A^, proteins were first pre-incubated at 16 °C for 30 min in the presence of E3 ligase reaction buffer. After that, the remaining reagents were added for the ubiquitylation reaction and incubated at 30 °C for the indicated time. For substrate ubiquitylation, *C. elegans* AHCY-1 was added as the substrate along with the other reagents and mixed with pre-incubated CHN-1 or pre-incubated CHN-1/UFD-2 and incubated at 30 °C for 90 mins. For performing the *in vitro* reaction in the presence of a chaperone, *C. elegans* 1 µM His-tagged HSP-1, His-tagged DAF-21, or other variants were pre-incubated with CHN-1 or CHN-1/UFD-2 at 16 °C for 30 min in the presence of 1x E3 ligase reaction buffer. After that, the remaining reagents were added for the reaction and incubated at 30 °C for 90 min. After the reaction, SDS-loading dye (Bio-rad, Cat#1610747), including β-mercaptoethanol (Sigma, Cat#M6250), was added to the entire reaction mix, and the samples were incubated at 95 °C for 5 min. Samples were run in 12% SDS-PAGE gels and blotted with an antibody against the protein of interest.

### E2 discharge assays

E2 discharging experimental protocol designed based on a modified method from Page et al., 2012. Discharging of increasing molar concentration (0.8, 1.6, 3.3, 5, 6.6 µM) of charged UBE2D1 (Boston Biochem, UBE2D1-Ub, Cat#E2-800) was performed at 30 °C for 40 min in ubiquitin conjugation reaction buffer (Boston Biochem, Cat#B-70). Similarly, a time-dependent assay was performed using 3.3 µM UBE2D1-Ub at 30 °C for different time points (5, 10, 20, 30 mins) with equimolar concentrations (1 µM) of CHN-1, His-tagged UFD-2^P951A^ and His-tagged HSP-1. The reaction was stopped by the addition of SDS-loading dye (Bio-Rad, Cat#1610747) without any reducing agent and incubation at 30 °C for 5 min. Samples were run in a 15% SDS-PAGE gel. For detecting the available UBE2D1-Ub in each condition, western blotting was performed using an anti-ubiquitin antibody. Normalized chemiluminescence intensity was obtained after maximum background subtraction from each lane.

### Western blotting and quantification

Protein samples in SDS-loading dye (reducing/non-reducing) were run in 12% or 15% acrylamide gels using running buffer (25 mM Tris, 190 mM Glycine, 0.1% SDS) at 120 volts (constant). The wet transfer was done at a constant 200 mA for 2 hr at room temperature using transfer buffer (25 mM Tris, 190 mM Glycine, 10% methanol, pH 8.3). Blots were then blocked with 5% skimmed milk in TBST (50 mM Tris, 150 mM NaCl, 0.1% Tween 20, pH 7.5) for 1 hr at room temperature. Blots were incubated with primary antibody prepared in 5% skimmed milk in TBST at 4 °C, overnight. The blots were then washed three times with TBST for 10 min each. Finally, the blots were incubated with secondary antibodies prepared in 5% skimmed milk in TBST for 1 hr at room temperature. The blots were imaged using a ChemiDoc™ Imaging System (Bio-Rad). All antibodies used in this study are listed in the resource table. We used Image Lab™ (version 6.0.0 build 25) software for blot quantification using Image Lab and graph plotted using GraphPad Prism 9. The bands appeared after probing with a particular antibody were marked in high sensitivity mode and quantified. Normalized chemiluminescence intensities were determined after maximum background subtraction from each lane.

### Enzyme-linked immunosorbent assay (ELISA)

2 µg/mL of UFD-2, His-tagged DAF-21, and His-tagged HSP-1 in coating buffer (100 mM NaHCO_3_, 32 mM Na_2_CO_3_, pH 9.2) were immobilized on Nunc-Immuno plates for ELISA (Thermo Fisher Scientific, Cat#44-2404) overnight at 4 °C. Blocking was performed with 2% BSA for 1 hr at 25 °C followed by washing with TBST (0.1% Tween 20). After incubation with increasing CHN-1 concentrations of for 1 hr at 16 °C, unbound CHN-1 was washed away by subsequent TBST washing steps. Interacting proteins were detected using an antibody against CHN-1 (1:5000 dilution, overnight 4 °C), followed by TBST washing and the addition of an HRP-conjugated secondary antibody. After the final wash, 100 µL of pnPP substrate (Alkaline Phosphatase Yellow-Sigma, Cat#P7998) was added in the dark. After 15 min, the reaction was stopped by adding 50 µL of 3M NaOH and the absorbance was measured at 450 nM.

### Modeling and Molecular Dynamics

CHN-1 model was generated by homology modeling using a Swiss model server (Waterhouse et al., 2018) with PDB ID 2F42 and 2C2L as the templates. The primary sequence of peptides used for docking on the CHN-1 dimer model was 628-640 HSP-1 (P09446) and 894-911 UFD-2 (Q09349). The protein and peptide complex structures were subjected to an energy minimization strategy using pmem.cuda (Goetz et al., 2012; Salomon-Ferrer et al., 2013) from AMBER18 (Case et al., 2018). We used tLeap binary (part of AMBER18) for solvating the structures in an octahedral TIP3P water box with a 15 Å distance from the structure surface to the box edges, and closeness parameter of 0.75 Å. The system was neutralized and solvated in a solution of 150 mM NaCl. AMBER ff14SB force field was used (Maier et al., 2015) and simulations were carried out by equilibrating the system for 1ns (NPT), at 1 atm, 300K, followed by 10ns NPT for non-bonded interaction. The particle mesh Ewald (PME) method was used to treat the long-range electrostatic interactions. Hydrogen bonds were constrained using SHAKE algorithm and integration time-step at 2 fs. (Ryckaert et al., 1977).

### Hydrogen deuterium exchange mass spectrometry (HDX-MS)

Prior to HDX-MS reactions, a complex of CHN-1 (3 mg/ml) and His-tagged UFD-2 (2 mg/ml) was formed by mixing the proteins in a 1:1 molar ratio followed by incubation at 25 °C temperature for 30 min. HDX-MS of CHN-1 and CHN-1 in complex with UFD-2 were performed at five time points during the incubation with deuterium (10 sec, 1 min, 5 min, 25 min, 2 hrs) in triplicate. 5 µl aliquots of protein were added to 45 µl of deuterated buffer (10 mM HEPES, 150 mM NaCl in 99.99% D_2_O; pH=8.0) at room temperature. The exchange reaction was quenched by moving the exchange aliquots to pre-cooled tubes (on ice) containing 10 µl of quenching buffer (2 M glycine, 4 M guanidine hydrochloride, 100 mM TCEP in 99.99% D_2_O, pH 2.3). After quenching, samples were frozen immediately in liquid nitrogen and kept at -80 °C until mass spectrometry measurement. Samples were thawed directly before measurement and injected manually onto the nano ACQUITY UPLC system equipped with HDX-MS Manager (Waters). Proteins were digested online on 2.1 mm x 20 mm columns with immobilized Nepenthesin-2 (AffiPro), for 1.5 min at 20 °C and eluted with 0.07% formic acid in water at a flow rate of 200 µl/min. Digested peptides were passed directly to the ACQUITY BEH C18 VanGuard pre-column from which they were eluted onto the reversed-phase ACQUITY UPLC BEH C18 column (Waters) using a 6–40% gradient of acetonitrile in 0.01% of formic acid at a flow rate of 90 μl/min at 0.5 °C. Samples were measured on the SYNAPTG2 HDX-MS instrument (Waters) in IMS mode. The instrument parameters for MS detection were as follows: ESI – positive mode; capillary voltage – 3 kV; sampling cone voltage – 35 V; extraction cone voltage – 3 V; source temperature – 80 °C; desolvation temperature -175 °C; and desolvation gas flow - 800 l/h. The CHN-1 peptide list was obtained using non-deuterated protein samples, processed as described above for HDX experiments, and measured in MSe mode. Peptides were identified using ProteinLynx Global Server Software (Waters). The HDX-MS experiment was analyzed using DynamX 3.0 (Waters) software. The PLGS peptide list was filtered by minimum intensity criteria – 3000 and minimal product per amino acid – 0.3. All MS spectra were inspected manually. Final data analysis was carried out using in-house HaDex software (Puchała et al., 2020). Differential deuterium exchange of residues was mapped to the model of CHN-1 generated using the 2C2L CHIP structure on the Swiss model server (https://swissmodel.expasy.org/).

### Surface plasmon resonance (SPR)

SPR-based interaction analysis was carried out at 25°C on a Biacore™ S200 instrument (GE Healthcare, Sweden). Recombinant purified His-tagged UFD-2 and His-tagged Ufd2p proteins were immobilized on NTA Biacore sensor Chips (Series S) at 20 μg/mL. Single-cycle kinetics studies were performed by passing increasing concentrations) (0, 100, 200, 500, 1000 and 2000 nM) of analyte M1 diUb conjugates (UbiQ, Cat#UbiQ-L01) in SPR buffer (10 mM HEPES, 150 mM NaCl, 0.05% Tween 20, 0.1% BSA, 50 µM EDTA, pH 8.0). The runs for both proteins were carried out under identical conditions. All injections were compiled in the same sensorgram with the response unit (RU) on Y-axis versus time (sec) on the X-axis.

### Preparation of *C. elegans* lysates and co-immunoprecipitation

Worms were grown at 20 °C. For protein extraction, worms were collected in M9 buffer and lysed using a lysis buffer (1M KCl, 1M Tris-HCL pH 8.2, 1M MgCl_2_, 0.07% NP-40, 0.7% Tween-20, 0.1% gelatine) with protease inhibitor (Roche, Cat# 11873580001) and in the presence of DUB inhibitor (Sigma-Aldrich, Cat#E3876). First, worms in lysis buffer were snap-frozen in liquid nitrogen. Next, the frozen samples were sonicated (40% amplitude, 5 cycles of 30 s pulses at 30 s intervals, Vibra-Cell™) on ice. Samples were centrifuged at 13,000 rpm for 15 min and the supernatants were collected. For co-immunoprecipitation, anti-DYKDDDDK (FLAG tag) magnetic beads (Pierce™ Anti-DYKDDDDK Magnetic Agarose, Cat#A36797) were used. 50 µl of anti-DYKDDDDK magnetic beads slurry were used for 200 µl of worm lysate. Lysate of CHN-1::FLAG-expressing worms was used as the experimental sample and wild-type (N2) worms were used as a negative control. Worm lysates were incubated with equilibrated magnetic beads at 4 °C for 1 and 2 hr for UFD-2 and AHCY-1 pull down, respectively. After the desired incubations, the beads were washed three times using washing buffer (PBS with 100 mM NaCl). Samples were eluted via the addition of SDS-loading dye (Bio-rad, Cat#1610747) containing β-mercaptoethanol (Sigma, Cat#M6250) and boiling for 5 min.

### RNA interference (RNAi)

RNAi was performed using the standard RNAi feeding method and RNAi clones (Kamath and Ahringer, 2003). For experiments, NGM plates supplemented with 1 mM IPTG and 25 µg/µL carbenicillin were seeded with HT115 *E. coli* expressing double-stranded RNA (dsRNA) against the gene of interest or, as a control, bacteria with the empty vector were used. Worms were placed on freshly prepared RNAi plates as age-synchronized L1 larvae.

### Proteomics

#### Protein digestion

For proteomic analysis, the following young adult strains were utilized: N2, *ufd-2(tm1380), chn-1(by155)* and *ufd-2(tm1380); chn-1(by155)*. For lysis, 4% SDS in 100 mM HEPES pH = 8.5 was used, and the protein concentrations were determined. 50 µg of protein was subjected for tryptic digestion. Proteins were reduced (10 mM TCEP) and alkylated (20 mM CAA) in the dark for 45 min at 45 °C. Samples were subjected to SP3-based digestion (Hughes et al., 2014). Washed SP3 beads (SP3 beads (Sera-Mag(TM) Magnetic Carboxylate Modified Particles (Hydrophobic), and Sera-Mag(TM) Magnetic Carboxylate Modified Particles (Hydrophilic)) were mixed equally, and 3 µL of beads were added to each sample. Acetonitrile was added to a final concentration of 50%, and the samples were washed twice using 70% ethanol (200 µL) on an in-house-made magnet. After an additional acetonitrile wash (200 µL), 5 µL of digestion solution (10 mM HEPES pH 8.5 containing 0.5 µg Trypsin (Sigma) and 0.5 µg LysC (Wako)) was added to each sample and incubated overnight at 37 °C. Peptides were cleaned on a magnet using 2 × 200 µL acetonitrile washes. Peptides were eluted in 10 µL of 5% DMSO in an ultrasonic bath for 10 min. Formic acid and acetonitrile were added to final concentrations of 2.5% and 2%, respectively. Samples were frozen until LC-MS/MS analysis. Liquid chromatography and mass spectrometry: LC-MS/MS instrumentation consisted of a nLC 1200 coupled to a nanoelectrospray source to a QExactive HF-x (Thermo Fisher Scientific) mass spectrometer. Peptide separation was performed on an in-house-packed column (75 µm inner diameter, 360 µm outer diameter), and the column temperature was maintained at 50 °C using a column oven (PRSO-V2). The LC buffer system consisted out of 0.1% formic acid (A) and 0.1% formic acid in 80% acetonitrile (B). Peptides were separated using a 90 min gradient applying a linear gradient for 70 min from 7 to 29 % B and then ramped to 65% B within 10 min, followed by a linear increase to 95% B within 5 min. 95% B was held for 5 min. Before each run, the column was re-equilibrated to 0%B. The mass spectrometer operated in a data-dependent acquisition mode targeting the top 22 peaks for collision-induced fragmentation and MS2 spectra acquisition. MS1 spectra were acquired in a scan range from 350 to 1650 m/z allowing a maximum injection time of 20 ms for an AGC target of 3e6. Spectra were acquired at a resolution of 60,000 (at 200 m/z). Ions were isolated in an isolation window of 1.3 m/z using an AGC target of 1e6 and a maximum injection time of 22ms. Spectra were acquired at a resolution of 15,000. The scan range for the MS2 spectra was set to 200–2000 m/z. The normalized collision energy was 28. Dynamic exclusion was set to 20 s. Data analysis: Acquired raw files were correlated to the Uniprot reference *C. elegans* proteome (downloaded: 06.2018) using MaxQuant (1.5.3.8) (Cox and Mann, 2008) and the implemented Andromeda search engine (Cox et al., 2011). Label-free quantification and matching between runs were enabled using default settings. Carbamidomethylation of cysteine residues was set as a fixed modification. Oxidation of methionine residues and acetylation of protein N-termini were defined as variable modifications. FDR was controlled using the implemented revert algorithm to 1% at the protein and the peptide-spectrum match (PSM). To identify significantly changed proteins, we performed a one-way analysis of variance (ANOVA) correcting for multiple testing using a permutation-based approach (FDR < 0.05, # permutations: 500).

### Lipidomics

The following young adult strains were utilized for lipidomic analysis: N2 (wild-type), *ufd-2(tm1380), chn-1(by155)*. Lipid extraction: Lipids from a homogenized sample comprising 15 000 worms were extracted using the Folch method as follows: 200 µL of methanol was added to each sample followed by 10 s of vortexing. Next, 500 µL of chloroform was added, followed by 10 s vortexing. This was followed by the addition of 200 µL of water to each sample to induce phase separation, following by vortexing for 20 s. The samples were then kept in the cold for 10 min. The samples were then centrifuged at 14.500 rpm for 10 min. The bottom layer was then pipetted out, and the solvent was dried under a stream of nitrogen. Prior to LC-MS analysis, the lipid extract was reconstituted in 200 µL of 1:1 isopropanol:methanol solution. LC-MS analysis: LC-MS analysis was performed as previously described (Nature Methods volume 14, pages 57–60 (2017)). Briefly, lipid extracts were separated on a Kinetex C18 2.1 × 100 mm, 2.6 µm column (Phenomonex, Aschaffenburg, De). Separation was achieved via gradient elution in a binary solvent, Vanquish UHPLC (Thermo Scientific, Bremen, DE). Mobile Phase A consisted of ACN:H_2_O (60:40), while mobile phase B consisted of IPA:ACN (90:10). For positive ionization, the mobile phases were modified with 10 mM ammonium formate and 0.1% formic acid, while for the negative ionization mode, the mobile phases were modified with 5 mM ammonium acetate and 0.1% acetic acid. A flow rate of 260 µL/min was used for separation, and the column and sample tray were held constant at 30 °C and 4 °C, respectively. 2 µL of each sample was injected onto the LC column. MS Instrumentation: MS analysis was performed on a Q-Exactive Plus Mass Spectrometer (Thermo Scientific, Bremen, DE) equipped with a heated electrospray ionization probe. In both the positive and negative ionization modes, the S-Lens RF level was set to 65, and the capillary temperature was set to 320 °C, and the sheath gas flow was set to 30 units and the auxiliary gas was set to 5 units. The spray voltage was set to 3.5 kV in the negative ionization mode and 4.5 kV in the positive ionization mode. In both modes, full scan mass spectra (scan range m/z 100–1500, R=35K) were acquired along with data-dependent (DDA) MS/MS spectra of the five most abundant ions. DDA MS/MS spectra were acquired using normalized collision energies of 30, 40, and 50 units (R= 17.5K and an isolation width = 1 m/z). The instrument was controlled using Xcalibur (version 4.0). Data analysis and lipid annotation: Progenesis Q1, version 2.0 (Non-Linear Dynamics, A Waters Company, Newcastle upon Tyne, UK) was used for peak picking and chromatographic alignment of all samples, with a pooled sample used as a reference. Lipids were annotated using the Progenesis Metascope Basic Lipids the LipidBlast databases with consideration made only of compounds that had MS/MS data. In both databases, the precursor ion tolerance was set to 10 ppm, and the fragmentation ion tolerance was set to 15 ppm. Putative lipid identifications were based on manual curation of database matches with fragmentation scores >10%.

### Fluorescent labeling of lipids in *C. elegans*

Lipid content in young adult worms was determined by RediStain™ WormDye Lipid Green (NemaMetrix, Cat#DYE9439) staining, according to the manufacturer’s protocol with incubation for 30 mins at room temperature with shaking. Working dye concentration: 1 µl of dye/200 µl of M9 buffer. Worms were protected from light, and several washes in M9 buffer were performed after staining. Immediately after that, imaging was performed on a Nikon SMZ25 microscope after immobilizing worms with tetramizole. Data analysis: Image processing was performed with ImageJ (Fiji) using Binary Mask and Particle Analysis Procedure with background signal subtraction. The graphs were plotted using GraphPad Prism 9.

### Yeast two-hybrid screening

Yeast two-hybrid screening was performed by Hybrigenics Services (http://www.hybrigenics-services.com). The coding sequence for *C. elegans* CHN-1 (NM_059380.5, aa 1–266) was PCR-amplified and cloned into pB27 as a C-terminal fusion to LexA (LexA-CHN-1). The construct was checked by sequencing the entire insert and used as a bait to screen a random-primed *C. elegans* mixed-stage cDNA library constructed into pP6. pB27 and pP6 were derived from the original pBTM116 (Vojtek and Hollenberg, 1995) and pGADGH (Bartel et al., 1993) plasmids, respectively. 61 million clones (6-fold the complexity of the library) were screened using a mating approach with YHGX13 (Y187 ade2-101::loxP-kanMX-loxP, mat□) and L40□Gal4 (mata) yeast strains as previously described (Fromont-Racine et al., 1997). 202 His+ colonies were selected on a medium lacking tryptophan, leucine, and histidine and supplemented with 50 mM 3-aminotriazole to prevent bait autoactivation. The prey fragments of the positive clones were amplified by PCR and sequenced at their 5’ and 3’ junctions. The resulting sequences were used to identify the corresponding interacting proteins in the GenBank database (NCBI) using a fully automated procedure. A confidence score (PBS, for Predicted Biological Score) was attributed to each interaction as previously described (Formstecher et al., 2005). The PBS relies on two different levels of analysis. First, a local score considers the redundancy and independence of prey fragments and the distribution of reading frames and stop codons in overlapping fragments. Second, a global score considers the interactions found in all of the screens performed by Hybrigenics using the same library. This global score represents the probability of interaction being nonspecific. The scores were divided into four categories for practical use, from A (highest confidence) to D (lowest confidence). A fifth category (E) flags explicit interactions involving highly connected prey domains previously found several times in screens performed on libraries derived from the same organism. Finally, several of these highly connected domains were confirmed as false positives and were tagged as F. PBS scores have been shown to positively correlate with the biological significance of interactions (Rain et al., 2001; Wojcik et al., 2002).

## Acknowledgments

We thank the *Caenorhabditis* Genetics Center (funded by the NIH National Center for Research Resources, P40 OD010440) for strains, Addgene for plasmids and Katarzyna Prokop and Marta Niklewicz for technical assistance. We thank Vishnu Balaji and Gabriele Stellbrink of the Hoppe laboratory for discussions and technical support and members of Pokrzywa laboratory for discussions and comments on the manuscript.

## Author contributions

The project was initiated in the laboratory of T.H. A.D., P.T., N.S., K.B., and W.P. designed and conducted experiments. U.S. and C.J.C. performed structural modeling and simulation analysis. R.M.G. performed the lipidomic analysis. N.A.S performed the bioinformatic analyses. H.N. and M.K. performed the proteomic analyses. K.D, D.C, M.D. performed the HDX-MS studies. W.P. (with input from M.N.) conceived the project and supervised the study. W.P. (with input from M.N.) secured the funding. W.P. and A.D. wrote the manuscript with input from M.N, T.H and C.J.C. The authors declare no competing financial interests.

**TABLE S1.**
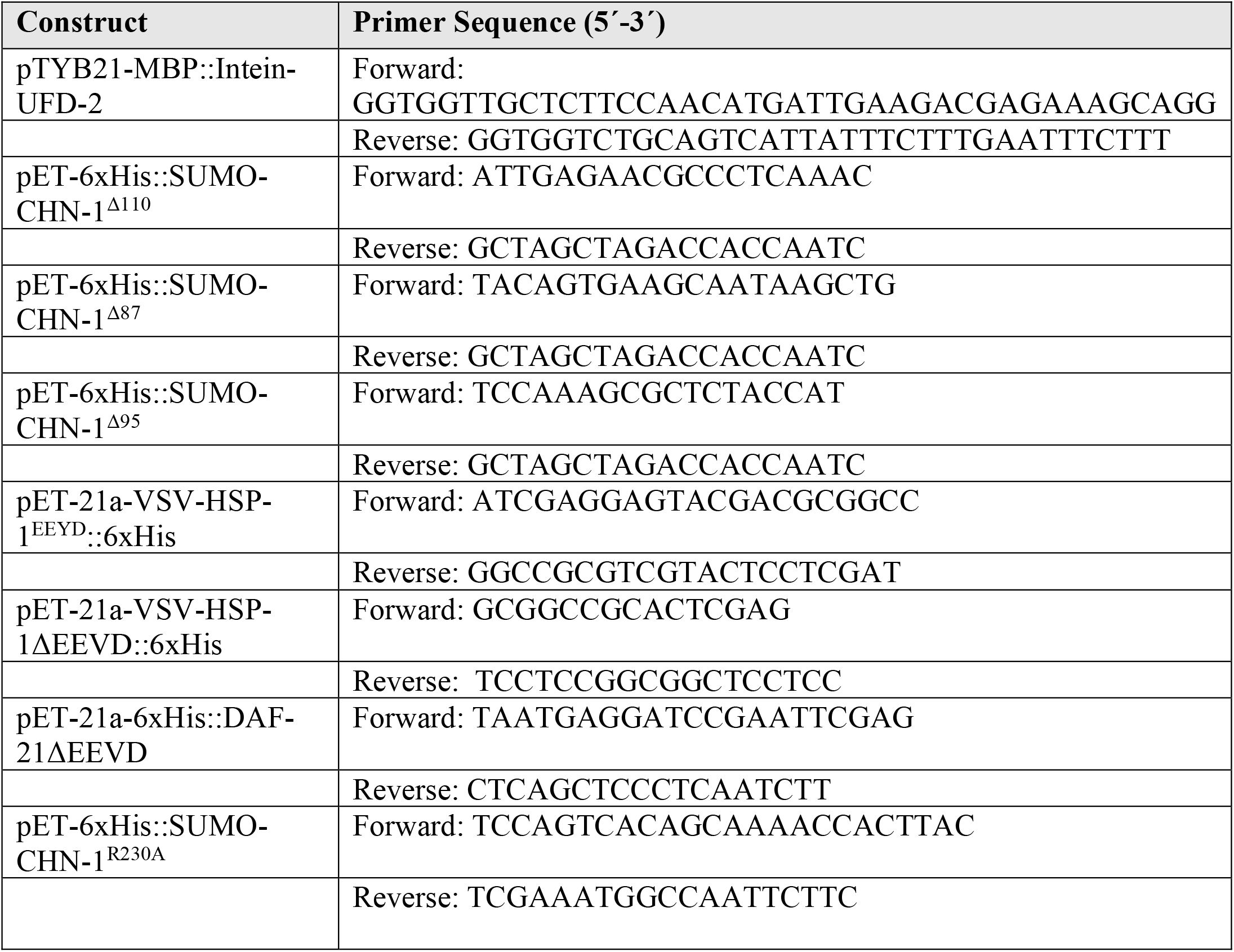
List of constructs and oligonucleotides used to generate them.

## KEY RESOURCES TABLE

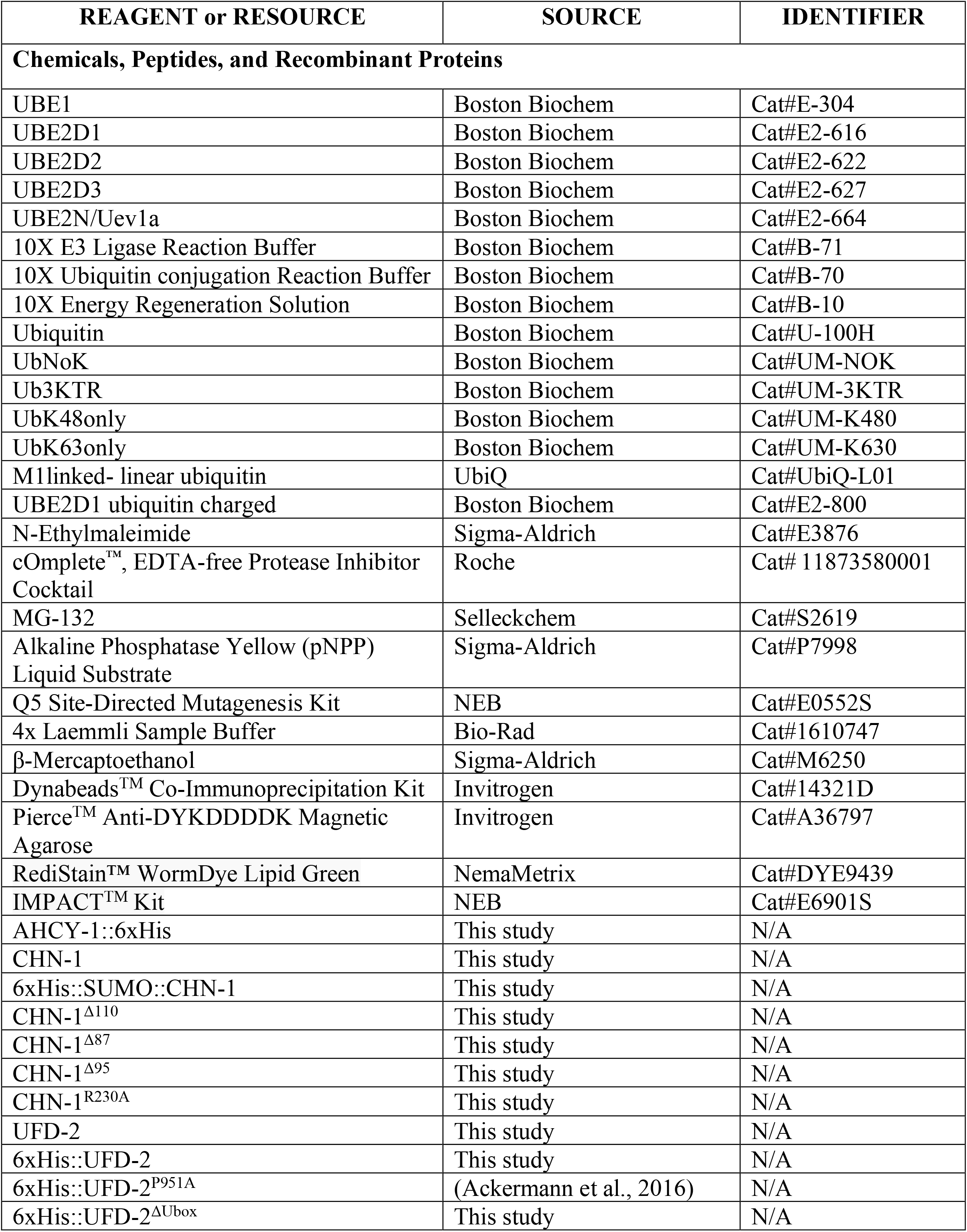
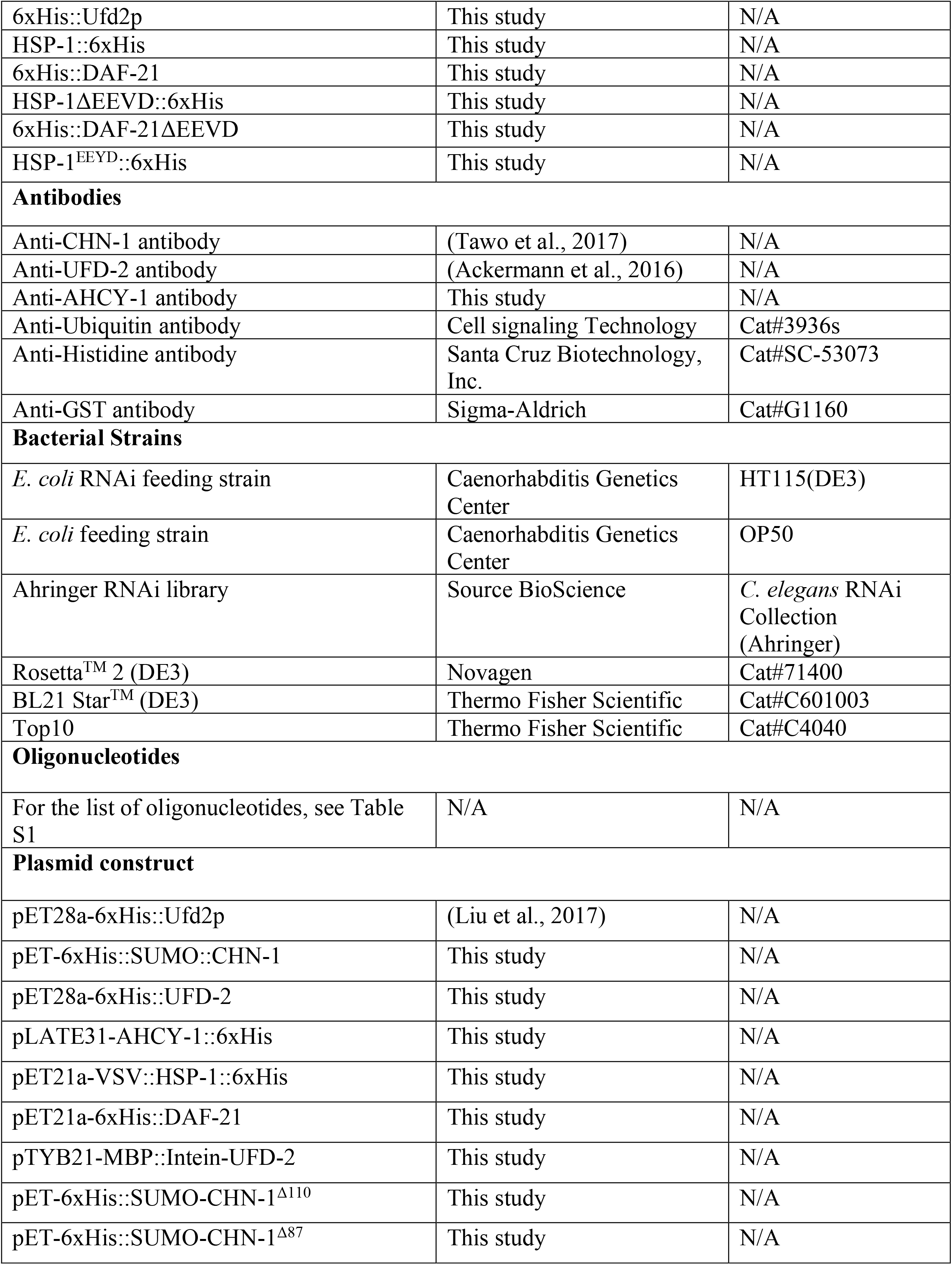
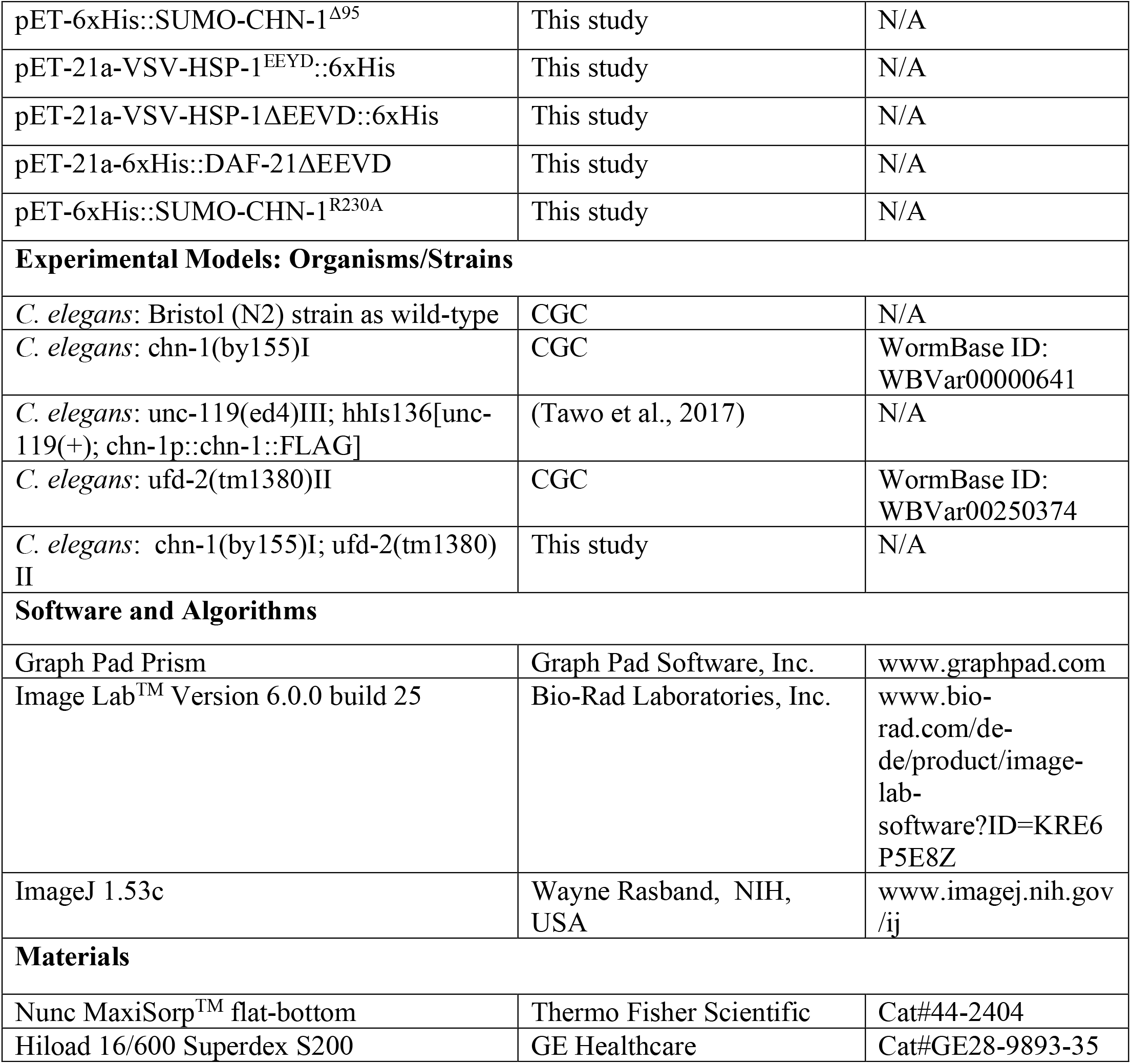

