## Supplemental Figure 1 for "Heterotypic Assembly Mechanism Regulates CHIP E3 Ligase Activity"

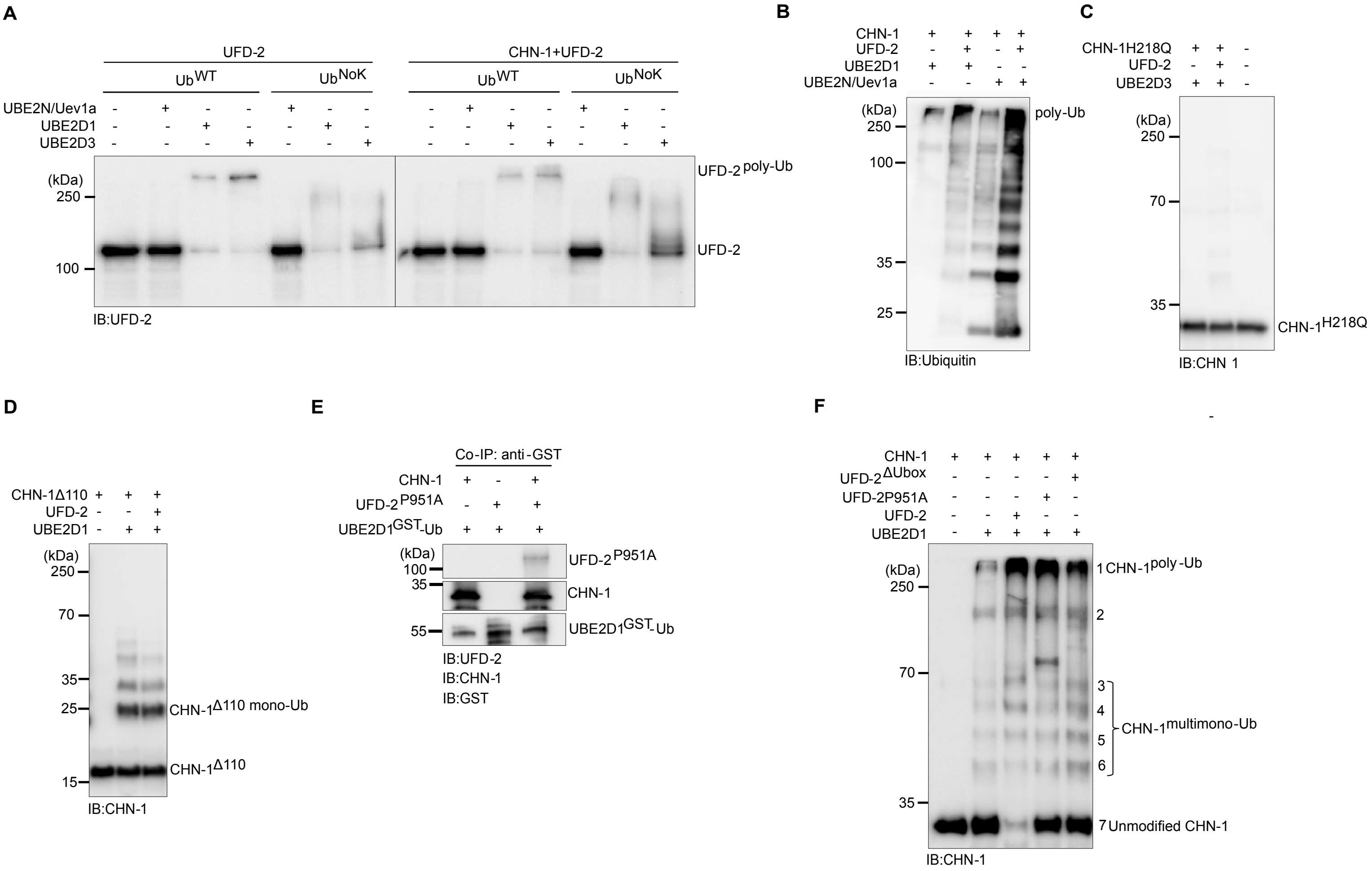


| | Band No. | CHN-1 -E2<br>Band % | CHN-1 +E2<br>Band % | UFD-2 CHN-1 +E2<br>Band % | UFD-2$^{P951A}$ CHN-1 +E2<br>Band % | UFD-2$^{\Delta Ubox}$ CHN-1 +E2<br>Band % |
|---|---|---|---|---|---|---|
| CHN-1$^{poly-Ub}$ | 1 | | 4.4 | 64.5 | 37 .8 | 27 .5 |
| | 2 | | 7.0 | 6.4 | 1.8 | 0.6 |
| CHN-1$^{multimono-Ub}$ | 3 | | 0.9 | 4.1 | 1.1 | 0.3 |
| | 4 | | 1.8 | 8.1 | 2.5 | 0.8 |
| | 5 | | 0.2 | 5.2 | 3.7 | 6.8 |
| | 6 | | 0.3 | 0.3 | 0.4 | 0.9 |
| Unmodified CHN-1 | 7 | 100 | 85.3 | 11.4 | 52 .6 | 63 .0 |
