## Supplementary figures and images for "Heterotypic Assembly Mechanism Regulates CHIP E3 Ligase Activity"

### Supplemental Figure 2

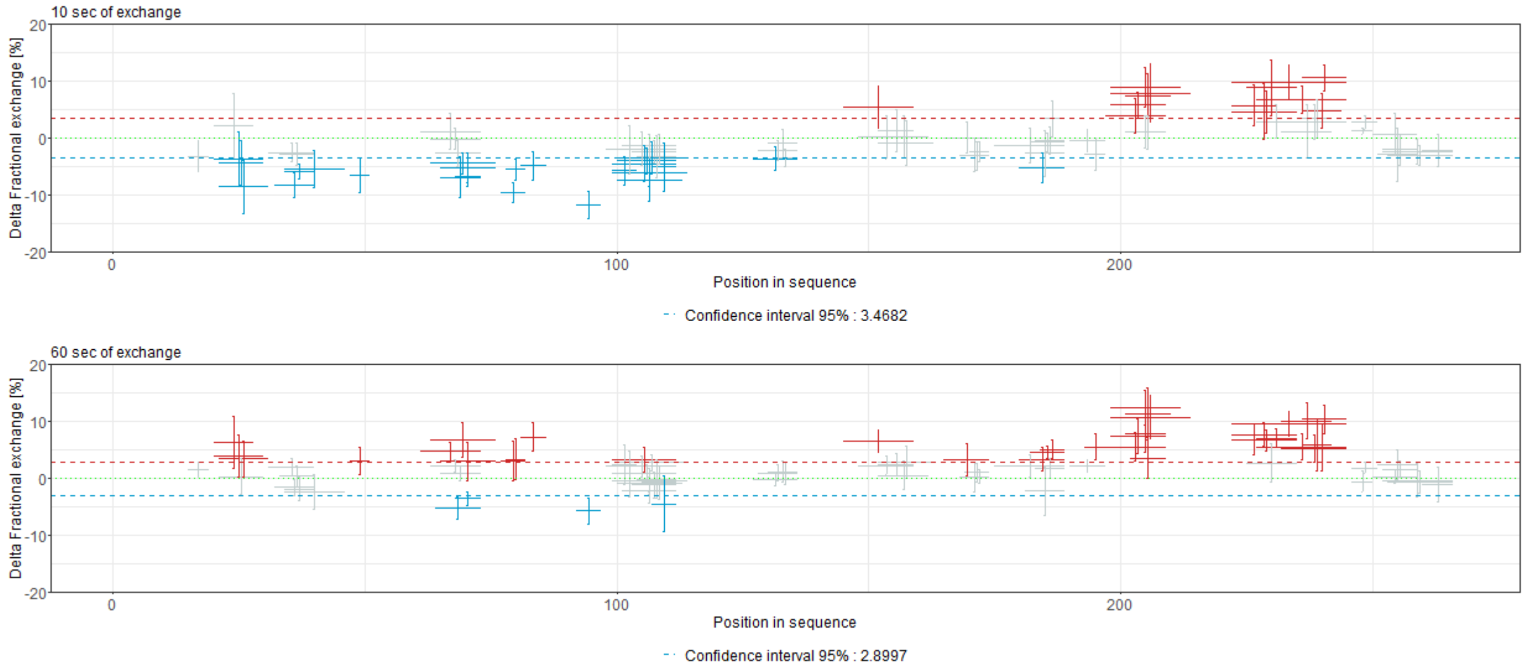

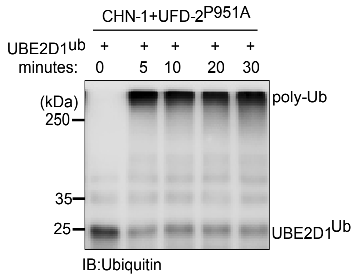

### Supplemental Figure 3

A

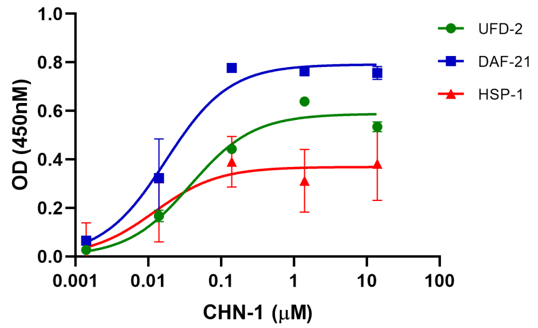


| HSP-1 $K_D$ | DAF-21 $K_D$ | UFD-2 $K_D$ |
|---|---|---|
| 12.17 nM | 17.44 nM | 37.82 nM |

B

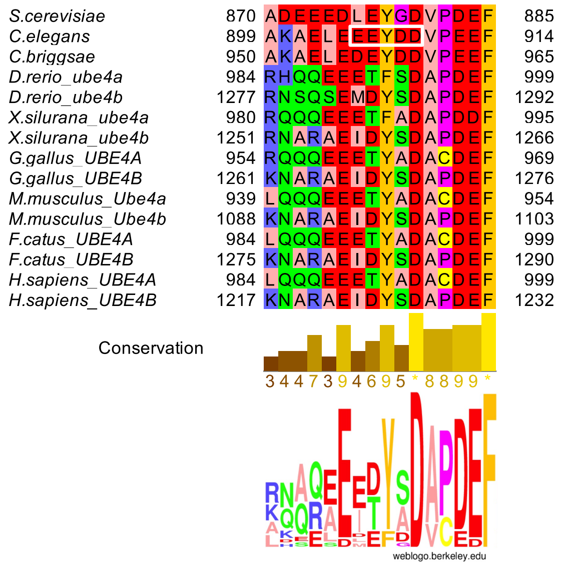


C

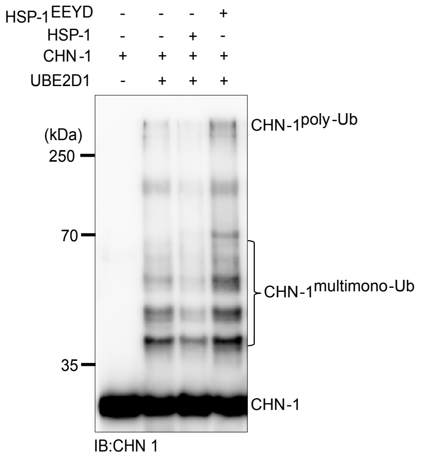


D

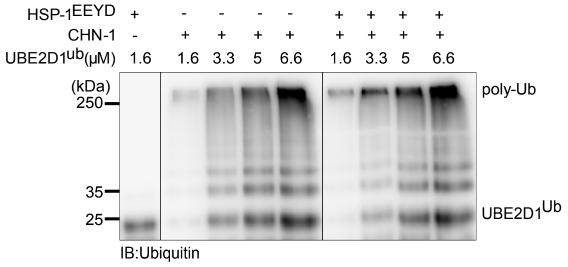


E

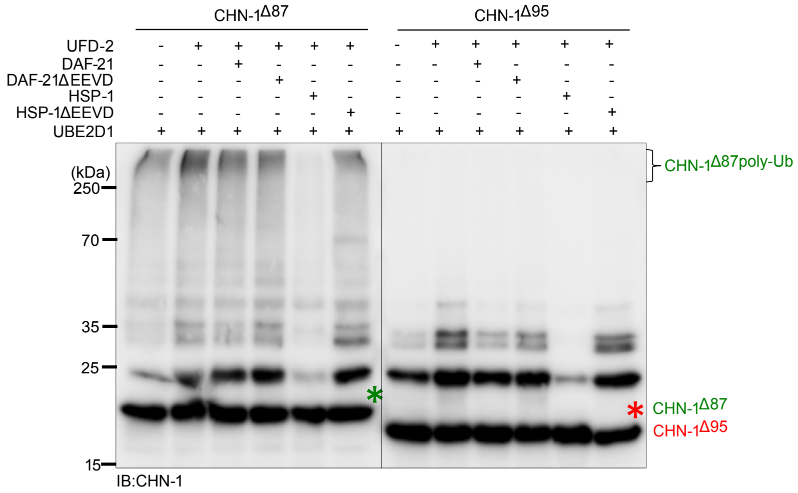


F

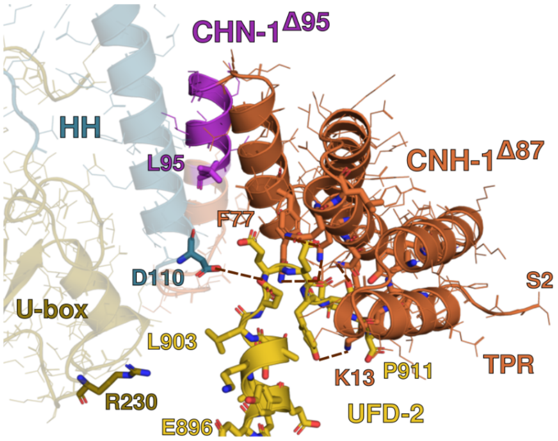


G

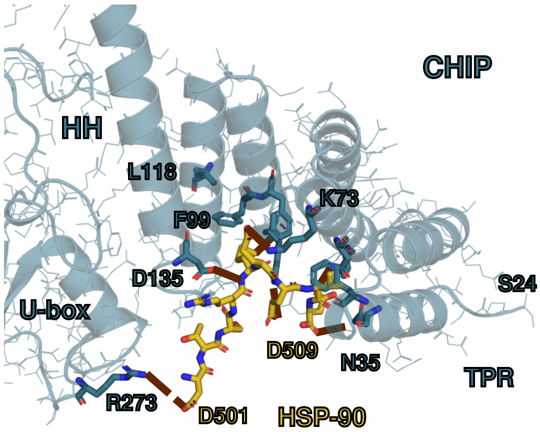

### Supplemental Figure 4

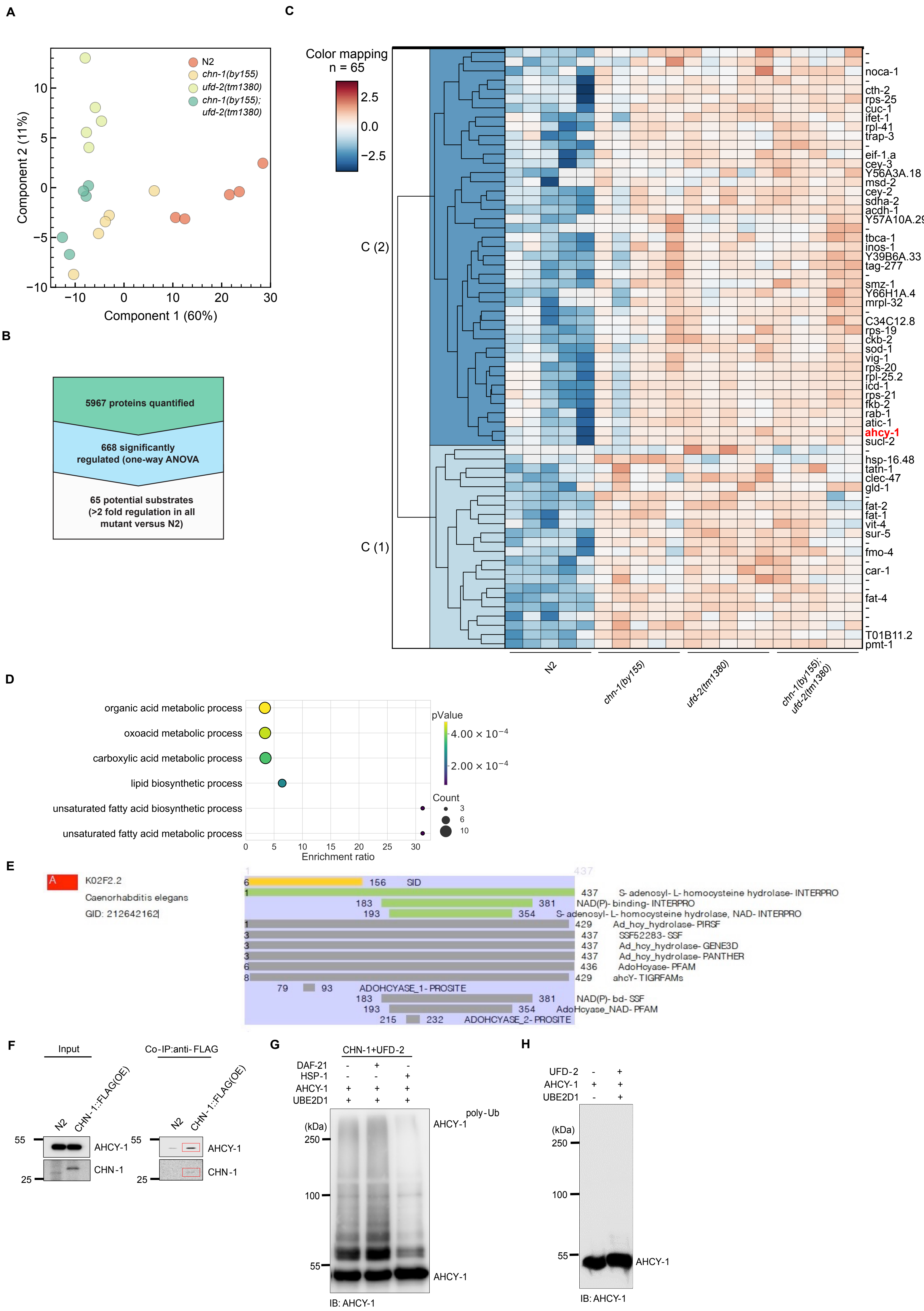
